# Unexpected post-release behaviours of reintroduced kākāpō

**DOI:** 10.64898/2026.09.24.753978

**Authors:** Andrew Digby, Scott W Forrest, Daryl Eason, Natasha Klappstein, Dan Howie, Craig Montgomerie, Janelle Ward, Kākāpō Recovery Team, Tāne Davis, Deidre Vercoe

## Abstract

Translocations are commonly used to reduce the risk of extinction of threatened species, and often to reintroduce animals to their former habitats. If extirpation occurred a long time ago, individuals may be placed into novel environments outside of conditions they have previously experienced, which can affect translocation success. To measure success of translocations, particularly those to novel habitats, it is important to monitor post-release spatial behaviour. In 2023, ten critically endangered kākāpō (*Strigops habroptilus*) were released to form a new population at the fenced mainland island of Sanctuary Mountain Maungatautari (hereafter Maungatautari) in the North Island of New Zealand, an area from which they were extirpated approximately a century earlier. The spatial behaviour and habitat selection of these kākāpō was analysed using dynamic space use and step selection functions with temporally dynamic responses applied to GPS logger data recorded over a seven month period. These showed three unexpected results: kākāpō were highly mobile and used larger areas compared to other sites; there was synchronicity of movement, with most individuals simultaneously moving toward the perimeter fence in spring; and when near the fence kākāpō selected for areas of pasture and exotic vegetation more than anticipated. While direct comparisons are difficult due to varying data availability, this unexpected spatial behaviour and habitat selection suggests that kākāpō behavioural responses were different at Maungatautari compared to other sites. This may indicate that kākāpō released at Maungatautari had a longer exploratory stage than elsewhere; that there were behavioural effects influenced by conspecific density or the availability of novel and exotic vegetation; or that there were specific breeding habitat requirements not met at the site during the study period. These results highlight the importance of high-resolution spatial monitoring to evaluate the success of translocations, particularly those to novel environments.

## 1. Introduction

Translocations play a key role in threatened species conservation, serving to reduce risk of extinction by re-establishing species in former habitats where they have been extirpated, or by augmenting existing populations to improve resilience of existing populations to fragmentation, changing climatic conditions, small populations and loss of genetic diversity (Armstrong et al., 1995; Hunter-Ayad et al., 2021; Seddon, 2010; Jarvie et al., 2014; IUCN, 2013). However, many translocations (≈20%) fail due to problems such as insufficient founder numbers, unsuitable habitat, dispersal away from release site, stress responses during acclimatisation period, or maladapted individuals (Berger-Tal et al., 2020; Bubac et al., 2019; Armstrong et al., 2013; Bilby and Moseby, 2024; Miskelly and Powlesland, 2013). Yet in many cases the reason for translocation failure is unknown, largely due to insufficient monitoring of individuals following release (Bubac et al., 2019; Tarszisz et al., 2014; Wilson et al., 2024). Even assessing success rates is difficult due to sparse documentation or well-defined criteria (Sutherland et al., 2010; Bubac et al., 2019; Parker et al., 2023).

It is possible to select potential release sites and timing using methods such as species distribution models (SDMs; Hunter-Ayad et al. 2021), which identify current and expected patterns of habitat use (Crego et al., 2021; Stone et al., 2025). This initial step can help improve translocation outcomes by selecting suitable habitats, but subsequent monitoring of translocated individuals’ behaviour in these novel environments is also essential in assessing translocation success. Post-release monitoring provides survival information and early detection of mortality risks, which can be related to individual behaviour (Zhang et al., 2025; Delcourt et al., 2025). After translocation, animals usually disperse before settling, and so the prolonged monitoring of individuals is necessary to distinguish between the exploratory and residency stages in order to successfully measure acclimatisation and detect excessive dispersal (Picardi et al., 2021; Delcourt et al., 2025; Wicks et al., 2024). Post-release monitoring also allows assessment of habitat selection and territory establishment (Berigan et al., 2024; Stone et al., 2025), as well as measurement of landscape connectivity (the extent to which landscape enables the movement of individuals; Berigan et al. 2024). These analyses can be used to assess habitat suitability, which is particularly difficult to predict in reintroductions (Stone et al., 2025) and which is important to link to measures of reproductive output in the residency stage (Armstrong and Seddon, 2008). Understanding habitat selection and dispersal can be used to target conservation management practices to improve survival and breeding success of the translocated population (Stone et al., 2025). Sufficiently detailed post-release monitoring is therefore essential in measuring and influencing the success of translocations.

Accurate measurements of post-release behaviour of individuals has been made possible by advances in tracking technology and statistical modelling (Katzner and Arlettaz, 2020; Trappes, 2023). Global positioning system (GPS) sensors can track spatial behaviour with high temporal and spatial resolution (Delcourt et al., 2025), and are particularly useful for monitoring cryptic individuals in habitat which is difficult to access or move around in (Wicks et al., 2024). Coupled with environmental data, GPS locations enable detailed analysis of habitat preferences and behaviour (Stone et al., 2025), which can subsequently be used to predict translocation risks that may vary due to seasonal or environmental effects or due to differences in individual behaviour and personality (Katzner and Arlettaz, 2020; Forrest, 2021; Delcourt et al., 2025; Spiegel et al., 2017). Understanding these risks can then be used to target interventions and plan translocations, even down to the individual level (Wicks et al., 2024; Katzner and Arlettaz, 2020; Buechley et al., 2018). Translocations are an important tool in the conservation of the kākāpō (*Strigops habroptilus*), a critically endangered, nocturnal and flightless parrot endemic to Aotearoa/New Zealand. Formerly one of the most common species and widespread throughout mainland New Zealand (Boast et al., 2025; Powlesland et al., 2006), it was nearly extirpated by a combination of habitat loss and predation by humans and introduced mammals (Dussex et al., 2021; Bergner et al., 2016; Carpenter et al., 2023). Two remnant populations located in the late 1970s were being preyed on by mustelids and cats (Powlesland et al., 2006), so the 62 remaining individuals were translocated to island sanctuaries free from mammalian predators (Powlesland et al., 1995). The species is now thought to be extinct throughout its historic mainland range (Carpenter et al., 2023).

By the mid-1990s, there were just 51 kākāpō left and the species faced challenges from infrequent breeding, high rates of infertility and embryo death, low genetic diversity and disease. An intensive conservation programme (the Kākāpō Recovery Programme, KRP) included radio-and activity-tracking of all individuals to monitor survival, and the support of breeding attempts with supplementary feeding, artificial incubation, hand-rearing and artificial insemination (Elliott et al., 2001; Eason and Moorhouse, 2006; Merton and Merton, 2006; Clout and Merton, 1998; Clout, 2006; Fischer et al., 2025). These efforts led to a recovery of the population to approximately 250 by 2023, resulting in the offshore island refuges approaching capacity and the advent of a new challenge to find suitable predator-free habitat.

Recovery of kākāpō has been complicated by the species’ unusual ecology. It is the only lek breeding parrot, with males congregating to conduct acoustic displays at “track and bowl” systems (Merton et al., 1984; Powlesland et al., 1992; Morris, 1977; Kelman, 2017). Kākāpō only breed during years of mass-fruiting (“masting”) of certain tree species (Henry, 1903), such as the rimu (*Dacrydium cypressinum*), which masts every 2 – 4 years (Cockrem, 2006; Powlesland et al., 1992; Cottam et al., 2006). Mothers feed nestlings almost exclusively on rimu fruit (Cottam et al., 2006).

Outside of nesting periods, kākāpō are generalist foragers of a wide range of vegetative matter, including leaves, fruits, bark, roots, mosses and fungi (Best, 1984; Boast et al., 2023; Atkinson and Merton, 2006; Butler, 2006; Horrocks et al., 2008; Johnson, 1976). Kākāpō can travel large distances each night and can have breeding and non-breeding ranges several kilometres apart (Best and Powlesland, 1986). Their range sizes vary with the habitat availability and season, but typically are in the range 15 − 50 ha (Joyce, 2009; Elliott, 2006; Whitehead, 2007; Walsh et al., 2005; Best and Powlesland, 1986; Farrimond et al., 2006; Powlesland et al., 2006).

Although they are phylogenetically and behaviourally unique and subject to intensive conservation management, knowledge of kākāpō ecology and behaviour is mostly limited to information from coprolites (Horrocks et al., 2008; Boast et al., 2025, 2023) and from observations of the small, isolated populations at relict mainland sites and at island sanctuaries (Powlesland et al., 1995). There are very few accounts of kākāpō behaviour in their original habitats (Henry, 1903; Hill, 2018), and studies of kākāpō spatial behaviour are especially limited, particularly following translocation. Joyce (2009) conducted the largest study of dispersal behaviour to date, following a population of 13 males and 13 females released on Pearl Island, adjacent to Rakiura/Stewart Island in southern New Zealand. Individuals exhibited substantial movement during the dispersal period, particularly females, although movement distances varied considerably among birds. In a smaller study, Trinder (1998) showed that translocated kākāpō tended to have larger ranges than resident birds, although the latter were anchored by supplementary feeding. Joyce 2009 also assessed spatial breeding behaviour, finding that both sexes travelled large distances during breeding, but there were no differences in home range size between the sexes, or between breeders and non-breeders. In a study of kākāpō space use on nearby Whenua Hou/Codfish Island during breeding in 2005, White-head et al. (2011) found that non-breeding females had smaller home range sizes than breeding females. To date there have been no peer-reviewed studies of kākāpō spatial behaviour beyond these, all of which used radio telemetry to triangulate kākāpō locations, so had limited spatial and temporal resolution compared to that available from GPS.

To address the lack of available predator-free island sites, in 2023 ten kākāpō were translocated to a mainland fenced reserve, Sanctuary Mountain Maungatautari in the central North Island (Smuts-Kennedy and Parker, 2013). This represented the first reintroduction of kākāpō to the mainland and to an area where they had been extirpated for at least a century (Carpenter et al., 2023). Subfossil data analysis showed that kākāpō were once present in the area (Lentini et al., 2018), but for the contemporary population which originated from Rakiura/Stewart Island and Fiordland, this translocation was to a relatively novel habitat type. Compared with the cooler podocarp-hardwood forests of Rakiura which are rich in rimu, kāmahi (*Pterophylla racemosa*) and southern rātā (*Metrosideros umbellata*), Maungatautari supports a warmer broadleaf-podocarp forest dominated by a rich assemblage of species such as tawa (*Beilschmiedia tawa*), pukatea (*Laurelia novae-zelandiae*), miro (*Prumnopitys ferruginea*) and mangeao (*Litsea calicaris*), with scattered large rimu and northern rātā (*Metrosideros robusta*). The density of rimu at Maungatautari is significantly lower than on the southern kākāpō islands such as Whenua Hou (Hurley, 2017). The translocation was the first kākāpō transfer to a fenced sanctuary — a site type which has been key to New Zealand native bird conservation over the last 30 years (Innes et al., 2012; Bombaci et al., 2018), but which had not previously housed kākāpō due to their insufficient size. With the advent of larger sites such as Maungatautari, fenced sanctuaries could provide important sites for kākāpō until large mainland predator-free areas are available (Bombaci et al., 2018).

Our objectives were to monitor post-translocation outcomes by conducting the highest resolution analysis of kākāpō movement ecology to date, the first on the mainland and the first within a fenced sanctuary. Specifically, we aimed to estimate kākāpō space use and habitat selection following translocation to Maungatautari by applying dynamic space use (Forrest et al., 2024) and step selection functions (Fortin et al., 2005; Thurfjell et al., 2014; Avgar et al., 2016; Klappstein et al., 2024) to GPS data and environmental variables. With little previous study of post-release behaviour of kākāpō, particularly into unoccupied sites, this study was primarily used to gather information into kākāpō dispersal behaviour, habitat utilisation and interactions with a mammalian pest-resistant fence. This information is important in informing suitable areas for future kākāpō release, identifying factors which affect translocation success, and assessing the suitability of fenced sanctuaries for kākāpō.

## 2. Methods

Sanctuary Mountain Maungatautari is a scenic reserve on an extinct volcano which reaches 797 m above sea level, located in the central North Island of New Zealand (Smuts-Kennedy and Parker, 2013). It is encircled by a 47 km mammalian pest-resistant fence to form 3363 ha of habitat which is free of mammalian pests except for mice and occasional incursions by rats, mustelids and possums. The vegetation is mostly native broadleaf podocarp forest (see Introduction for vegetation information), but with significantly lower density of rimu than on the southern kākāpō (Hurley, 2017). *Kākāpō release* islands Kākāpō were translocated to Maungatautari from islands in southern New Zealand in two releases: four individuals in July 2023 and six in September 2023. The kākāpō transferred were all males, comprising seven juveniles (hatched in 2019) and three adults (hatched in 2008, 2014 and 2016). Female kākāpō are of greater value for population growth, so males of lower genetic value are used to test new sites.

The kākāpō were released at different locations within the reserve, but all on tracks within 2 km of the fence (Figure 1). Prior to the kākāpō release, the Maungatautari perimeter fence was modified with a 38 cm metal baffle added to the inside mesh to deter kākāpō from climbing it.

**Figure 1:**
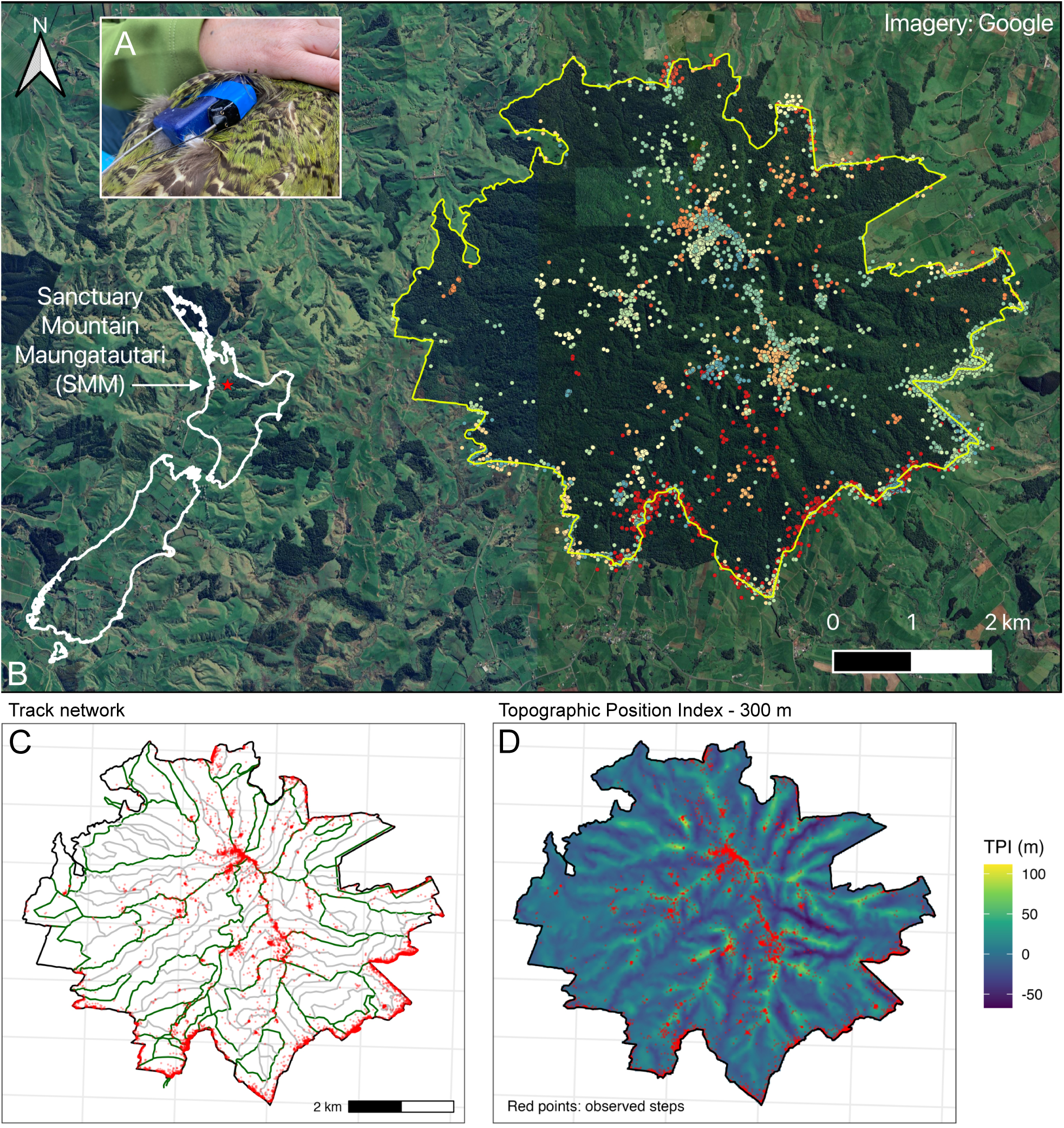
A) A Lotek PinPoint 240 GPS logger attached to a VHF transmitter on a kākāpō. B) Location of Maungatautari with respect to New Zealand, showing a satellite image of the study site and surrounding area. Each coloured point is the GPS location of a kākāpō, with different colours representing different individuals. C) The network of walking and maintenance tracks within Maungatautari, in relation to the GPS locations of all individuals, shown as red points, after points were clipped and reflected in the fence perimeter (see Methods). D) Topographic Position Index (TPI) of all cells within Maungatautari, where the TPI is the elevation difference between the focal cell and the mean of all cells within a 300 m circular radius, which quantifies ridges and valleys.

Despite this baffle being effective in trials (New Zealand Department of Conservation; NZDOC, internal report), kākāpō were detected outside the fence soon after the second release in September 2023. Subsequent trials indicated that this was achieved by climbing on vegetation which overhung the fence (NZDOC, internal report). From October 2023 to April 2024 there were 15 fence crossings involving six individuals, each of which crossed two to four times. Kākāpō were initially translocated back inside the fence following these movements, but by April 2024 seven of the ten kākāpō had been transferred back to southern islands to reduce the monitoring workload.

### GPS deployment

Each kākāpō was fitted with a ‘smart’ VHF transmitter (Lotek NZ Ltd, Havelock North, New Zealand and WildTech Ltd, Hastings, New Zealand) mounted on a ‘backpack’ harness, enabling VHF tracking and relay of activity information to the internet via a LoRA data network (designed and manufactured by NZDOC; Figure 1). These transmitters did not record positional information, so a separate GPS logger (Lotek PinPoint 240, 8 g, up to 3000 fixes per charge) was attached to the VHF transmitter. These were a “store-on-board” design, so had to be physically retrieved from the kākāpō to recharge and download the data. They were attached to the side of the VHF transmitter using 3M VHB double-sided tape (3M Ltd, Auckland, New Zealand), with electrical tape then added for extra security (Figure 1). These tags had been deployed in the same configuration on kākāpō at other sites since 2018.

The data used in this study were collected as part of routine kākāpō conservation management conducted by NZDOC as required by the New Zealand Conservation Act (1987), and so this study was exempt from the separate requirement of animal ethics approval under NZDOC’s obligations to the New Zealand Animal Welfare Act (1999).

The PinPoint loggers were programmed to attempt fixes every 30 minutes at night when kākāpō were active and once during the day when they were roosting. More fixes were not attempted during the day because kākāpō often roost in holes with poor satellite reception, which would lead to greater battery drain. The PinPoint tags operated on clock time rather than solar time, but were scheduled to attempt fixes during civil twilight and night (Figure S1). The Lotek ‘Swift’ fix algorithm was used to enable shorter fix times and longer battery life (Forrest et al., 2022). The GPS loggers were programmed to collect fixes over the mean estimated battery life, which was approximately three months (Figure S1). During January and February 2024 the PinPoint loggers on four individual kākāpō were programmed with more frequent five minute fixes to help identify fence crossing locations.

From January 2024 the Lotek PinPoint GPS loggers were replaced with a remote-download tag (Ornitrack-9 4G GPS, Ornitela, Vilnius, Lithuania) in order to be able to quickly detect kākāpō moving beyond the fence. The Ornitela tags had a less frequent fix schedule of 4–6 times per night, so the analysis described here was restricted to the “tracking period” of 19 July 2023 to 24 Feb 2024 when at least three kākāpō had continuous Lotek GPS data. Not all of the ten kākāpō had full GPS data coverage during this study period due to their later arrival and earlier departure from the sanctuary (Figure S1).

### GPS data and location filtering

Location errors were removed using speed filtering rather than the number of satellites or the horizontal dilution of precision (HDOP), since this ensures the removal of unrealistic locations and the retention of accurate locations that may be otherwise discarded (Forrest et al., 2022; Shimada et al., 2012). See Section B.1 for details of the speed filtering. The speed-filtered GPS data were further cleaned to remove data when individuals were known to have been outside of the fence, as determined by radio telemetry tracking and visual inspection of GPS tracks.

The GPS fix intervals were irregular for several reasons: i) fix attempt rates were different between day and night (Figure S1); ii) there were some periods and individuals with 5 minute fix attempts; and iii) the night-time fixes were frequently unsuccessful. A different resampling approach was used for the dynamic space use and the step selection analyses due to their differing statistical models. Because the occurrence distributions used in the dynamic space use analysis are sensitive to the fix schedule (Fleming and Calabrese, 2017; Alston et al., 2026), these locations were down-sampled to remove all fix intervals shorter than 30 minutes (Figure S3). This was achieved by specifying an asymmetric time tolerance in the track resampling, so that fix intervals of less than 30 minutes were downsampled to a minimum of 30 minutes and fix intervals longer than 30 minutes were retained to minimise data loss. For the step selection functions, all fix intervals were retained, and those that were less than 3 hours were grouped together using a “steps-by-burst” approach using the R package amt (Signer et al., 2019). As fix intervals were now a minimum of 5 minutes and a maximum of 3 hours, the irregular time intervals were accommodated within the movement kernel (see below).

Kākāpō spent large amounts of time next to the fence, meaning that measurement error typical to GPS devices resulted in reported locations that were outside of the fence when the kākāpō were known to be inside it (Figures S4 and S5). Assuming that GPS errors are considered to be radially symmetric (equally likely in any direction; Hurford 2009; Fleming et al. 2020), rather than discarding these locations, we “reflected” any locations that were beyond the fence by their distance outside of the fence to that distance inside the fence — i.e. a location that was measured as 15 metres outside the fence was placed 15 metres inside the fence on the reverse bearing. Any locations that were more than 200 m outside of the fence were removed from the data, as were any that were still outside the fence after reflection, due to being reflected across a narrow “peninsula” of fenced area. Before reflection, 18% of the SSF dataset (3006 of 16454) was outside of the fence but just 0.6% of the total (99) were discarded following the reflection process. We provide more information about the reflection process in the Supporting Information.

### Dynamic space use

The space used by the kākāpō during the tracking period was dynamic and changed continuously, likely reflecting their exploration of the novel environment. To assess the areas that were used and how they changed through time, temporal changes in kākāpō spatial behaviour at Maungatautari were analysed using temporally overlapping occurrence distributions (ODs), referred to as dynamic space use (Forrest et al., 2024). Forrest et al. (2024) used a dynamic space use approach to estimate temporal variability in risk to kākā (*Nestor meridionalis*) by their movements inside and outside a fenced sanctuary, and we adopt the same sweeping window framework here to assess changes to kākāpō space use within the fenced area at Maungatautari. ODs differ from long-term home ranges (range distributions) in that they estimate where an animal may have travelled during a defined period, measured as the probability of finding an animal at a particular location during a specified time, rather than estimating space use over the longer term (i.e. a home range; Alston et al. 2026). For the dynamic space use sweeping window (i.e., a one-dimensional convolution), we used a window width of 30 days that was moved across the full tracking period in steps of five days to select a series of temporally overlapping GPS fixes for each individual (refer to Forrest et al. 2024 for an illustration). The OD for each time interval was calculated using Kriged estimates implemented in the R package ctmm (Fleming et al., 2016), which accommodates a range of continuous-time movement models (Alston et al., 2026; Horne et al., 2007; Kranstauber et al., 2012). The fitted movement model was anisotropic Ornstein-Uhlenbeck Foraging (OUF), which was used for all individuals and time periods (Fleming et al., 2016).

As each individual was resampled to have the same GPS sampling approach, we compared relative changes in space use area and related the distance of the OD centroid to the sanctuary fence. The relative change in the space use area over the tracking period was calculated by comparing the 95th percentile of the OD area (OD95) for each time period, which is the smallest area that contains 95% of the probability density, to the mean of all ODs for that individual. Due to the exploratory and transient space use behaviour of the kākāpō, we did not estimate their long-term home ranges, but to quantify the overall area that each individual used we calculated 95% minimum convex polygons (MCP95) for the full trajectory of each individual.

### Step selection functions

An SSF assumes that we have a track of animal locations {***s***_1_, ***s***_2_,…,***s****_T_* } collected at regular time intervals, and defines the probability of a movement step ending at location ***s****_t_*_+1_ given the previous locations ***s***_1:_*_t_* = {***s***_1_, ***s***_2_,…,***s****_t_* } as

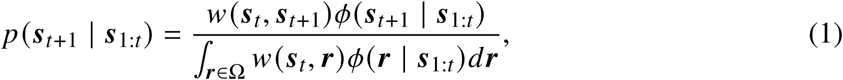

where *w* is a function describing habitat selection, *φ* is a movement kernel, and Ω is the spatial domain of integration. A key aspect of SSFs is that the integral in Equation 1 cannot be computed analytically and is approximated via numerical integration (described below, and in Michelot et al., 2024).

### Movement kernel

To approximate the SSF likelihood, we paired each observed location with *N* = 20 random points to form a stratum at each time. Unlike SSFs that estimate movement jointly with habitat selection (e.g., Forester et al., 2009; Avgar et al., 2016), we sampled random points from empirical distributions of step lengths and turning angles (as in Fortin et al., 2005). Michelot et al. (2024) shows that this approach can be viewed as a form of Monte Carlo integration, where the movement kernel is assumed to be known prior to model fitting (i.e., where *φ* is defined by the random points). To account for irregular time intervals arising from missing GPS fixes, we grouped observed steps into time bins based on the elapsed time between successive locations. Let Δ*t_t_* denote the time interval (in minutes) between locations *t* and *t* + 1, and let *b_t_* denote the corresponding time-bin index, defined as

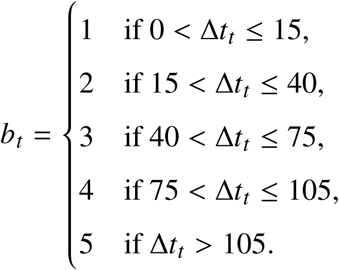

For each location ***s****_t_*, random points {***r***_1_*_t_*, ***r***_2_*_t_*,…,***r****_Nt_* } were then sampled from a movement kernel specific to the corresponding time bin, yielding the approximate SSF as,

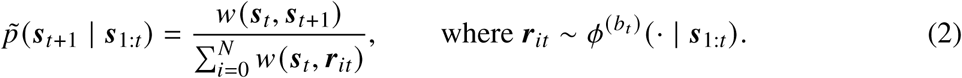

By specifying the distribution of random points *φ*^(bt)^ (· | ***s***_1:_*_t_*) to vary with time bin *b_t_*, this is analogous to including interactions on the movement kernel parameters. Similar methods have been shown to result in negligible bias in SSFs with movement covariates (Hofmann et al., 2024), but we chose empirical sampling to reduce the number of parameters to estimate, as we wished to prioritise complexity in the habitat selection function. To ensure equal numbers of random steps within each strata, the random steps were initially oversampled, with 40 drawn for every observed step. Any random steps that fell outside of the fence were removed (see *GPS data and location filtering*), since they were not considered available steps. From the random steps remaining, we randomly selected 20 which were used for model fitting. Any strata with fewer than 20 random steps or no observed step after fence reflection were discarded.

### Habitat selection function

The SSF was defined as a generalised additive model (GAM) with a Cox PH distribution using R package mgcv (Wood, 2011), which allows for flexible and non-linear responses to environmental covariates, including non-linear interactions with time (Klappstein et al., 2024). Specifically, following Klappstein et al. (2024), we defined the habitat selection function as

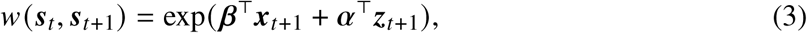

where ***x****_t_*_+1_ are covariates of the fixed effects ***β*** and ***z****_t_*_+1_ are the covariates of random effects ***α***. This formulation allows any covariate effect to be modelled as either a log-linear term, non-linear function, or with random effects. All model parameters were estimated using restricted maximum likelihood (REML; Klappstein et al. 2024).

The landscape variables used for input to the SSF were selected from spatial variables which were considered ecologically relevant. Understanding the interaction of kākāpō with the fence was a primary motivation of this study, so we included the (log-transformed) distance of each location to the fence at its closest point (Cowan et al., 2025). We included the natural log of distance to the nearest walking or maintenance track, as kākāpō are known to move using tracks (Figure 1C). To account for the topography of the sanctuary we used the topographic position index (TPI), which compares the elevation of a pixel to the mean of surrounding pixels within a specified radius. TPI quantifies ridges and valleys, which are expected to be relevant to kākāpō as ridges are often chosen as lek locations (Figure 1D). We included TPI with a radius of 300 m (Figure 1D), which was selected via model fitting (details in Supporting Information).

We also included vegetation cover, which originally had 10 classes (Table S2). To simplify the analysis, the two pasture categories (high-producing and low-producing), the two scrub categories (mixed native and unspecified scrub), and the two podocarp categories (mid altitude podocarp-hardwood forest and lowland podocarp-hardwood forest) were each merged into single pasture, scrub and podocarp categories respectively (Figure S7). The “pasture” category contained mostly pasture grasses, but also included a variety of introduced plants, particularly close to the fence. If any observed or random steps had missing vegetation values due to incompleteness of the vegetation map, then all steps in that strata were removed. GIS shape files for the fence line, tracks, elevation and vegetation cover were obtained from Maungatautari management data.

All covariates were checked for multicollinearity using Spearman and Pearson correlation coefficients. Elevation was highly correlated with the log of fence distance (Pearson *r* = 0.82), so elevation was removed as the distance to fence was of most interest. All remaining variables had variance inflation factors (VIF) of less than two. The final set of variables included in the SSF model were the distance to fence (log), distance to the nearest track (log), TPI at 300 m, and a vegetation layer with categories of podocarp, pasture and scrub. The continuous variables were standardised to mean of 0 and standard deviation of 1 to improve numerical stability and aid in interpretation.

### SSF model selection and validation

Linear terms were included for the (log) distance to nearest track, TPI, and categorical vegetation type, with a smooth interaction term (thin plate spline) included for the (log) distance to fence across time. To account for pseudo-replication, random effects were included for individual kākāpō. We included a random slope (bs =‘re’) for (log) track distance, TPI and vegetation, and a factor smooth interaction (bs = ‘fs’) for (log) distance to fence across time, which fitted individual-specific smooths of time for the effect of log distance to fence that were penalised towards the population smooth (Pedersen et al., 2019). An alternative model was also fitted with random slopes (bs = ‘re’) for the (log) distance to fence term at the individual-by-day level, instead of a factor smooth. The two models were compared with AIC scores and using cross-validation (see below; Fortin et al. 2009). Differences among vegetation classes were evaluated using post-hoc Wald contrasts of model coefficients.

Model validation is difficult for SSFs since residuals plots cannot be easily interpreted, particularly when using smooth terms (Klappstein et al., 2024). Instead, a cross-validation approach was used to assess the performance of the SSF models. We conducted k-fold cross-validation, following the procedure of Fortin et al. (2009) and adapted from Hofmann et al. (2023) and Pike et al. (2026). For each model, this involved splitting the data into training and testing datasets, where 80% was used to fit the model, and the remaining 20% was used for testing the model (selected randomly, preserving strata). For testing, step selection probability weights for the observed and randomly sampled steps were calculated using the estimated parameters of the full model using the predict.gam function from the mgcv package (Wood, 2011). The weights of the steps were ranked, where a rank of 1 was the lowest weight (least likely step), and a rank of 21 had the highest weight (most likely step). We extracted the rank of each observed step, where we would expect that a good model would place high weight (i.e. high ranks) on the observed steps, and calculated the Pearson correlation coefficient (*r_p_*). Values of *r_p_* can range from –1 (negative linear correlation) to 1 (positive linear correlation), with 0 indicating a model performs as well as random chance (providing the movement kernel is representative of the observed data; Michelot et al. 2024). As a control, we also randomly selected a single step per stratum and also calculated *r_p_*. We repeated the data splitting, model fitting and cross-validation process 100 times for each of the two models and calculated the mean and upper and lower 95% confidence intervals of *r_p_* for the observed and random selection (see Supporting Information; Figures S9, S10).

### Selection for pasture near fence

In order to determine why kākāpō might be frequenting the area near the fence, a *post-hoc*resource selection analysis was undertaken to assess if particular vegetation types were being used in the vicinity of the fence. Since “pasture” (which also included other exotic vegetation) was the most common vegetation type close to the fence, we assessed use of pasture compared to non-pasture areas. For each kākāpō, 1000 random points were selected within the intersection of the 95% minimum convex polygon (MCP95) for that individual and a 100 m buffer inside the fence. The proportion of random and observed points at each vegetation type within this area were then compared using a Chi-squared test.

## 3. Results

### GPS data and location filtering

From 36,250 fix attempts for all individuals there were 18,891 successful fixes, which were reduced to 17,014 after filtering for date and speed, resulting in a mean fix success rate of 45% over a combined deployment length of 1,352 kākāpō-days (Table S1). After removing fixes for fence breaches, downsampling the fix intervals and reflecting across the fence, the total data size for the dynamic space use analysis was 12,625 locations. After removing fence breach locations, reflecting across the fence, track bursting and resampling, random step sampling, and downsampling, the data set input to the SSF models contained 10,918 locations. Due to their exploratory and dynamic behaviour, the individual MCP95 area ranged from 712 ha to 2916 ha, with a mean across all individuals of 1,761 ha (Figure S12).

### Dynamic space use

Many of the ten kākāpō utilised large areas of the sanctuary and exhibited considerable movement during the study period, as indicated by the occurrence distributions (Figure 2 and Figures S13–S22). There was a synchronised change among the kākāpō in the relative area (compared to the mean) of the occurrence distribution over time. Most kākāpō utilised large relative areas from release until late October 2023, then reduced to much smaller areas throughout November and December, before increasing space use in late December and early January (Figure 2). The distance of each OD to the fence followed a similar temporal trend to space use area, with most individuals using space far from the fence at the start of the tracking period, near the release locations, and then moved towards the fence as the tracking period progressed.

**Figure 2:**
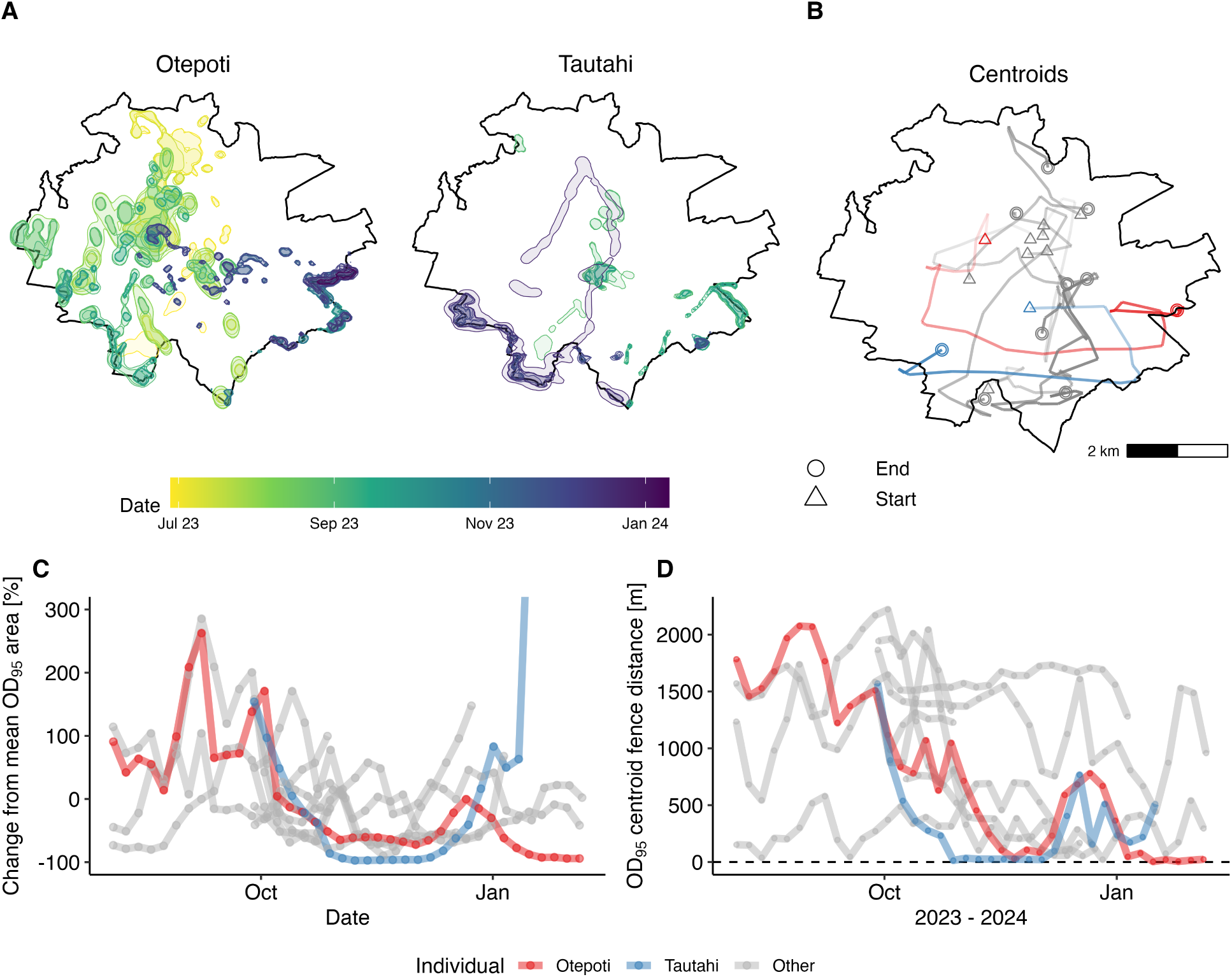
(A) 95% occurrence distributions (OD95; from Krige estimator) for two representative individuals, Otepoti and Tautahi, with colours denoting the time across the tracking period. (B) the change in OD95 centroid locations over time, with triangles denoting the centroid for the first time period, and circles that for the last. Line opacity increases with time, and individuals Otepoti (red) and Tautahi (blue) are highlighted. (C) the difference in OD95 area from the mean for each individual, and (D) the distance of the OD95 centroid from the fence, with the individuals in panel (A) highlighted. In panel (C) the vertical axis is truncated for clarity, omitting the final OD95 area difference for Tautahi of 581%.

### Step selection functions

Model performance parameters showed that the model with a factor smooth for fence distance was better fit than that with a random slope (Δ*AIC* = 37.6). Cross-validation showed that both models performed equally well, with a correlation coefficient *r_p_* = 0.744 for both the factor smooth model and the random slope models (Figures S9–S11). Based on these combined results, the factor smooth model was selected as the final model used for inference.

The overall effect for all kākāpō showed a significant decrease in the (log) distance to fence over the tracking period, indicating that the kākāpō moved towards the fence following release (Figure 3). However, there were marked differences in the individual responses, particularly as some individuals moved away from the fence towards the end of the monitoring period (Figure 3 and Figure 2D). The SSF model also showed that across all time periods and across the entire site, the kākāpō tended to select for habitat closer to tracks, and that there was a significant selection for ridges (higher TPI; Figure 3). Selection was greater in pasture than in scrub (*β* = 0.48 ± 0.16 SE, *p* = 0.003) and a direct contrast indicated that selection for pasture was also greater than for podocarp-hardwood vegetation (*β* = 0.34 ± 0.15 SE, *z* = 2.24, *p* = 0.02). Selection for podocarp-hardwood vegetation did not differ from scrub (*β* = 0.14 ± 0.14 SE, *p* = 0.315).

**Figure 3:**
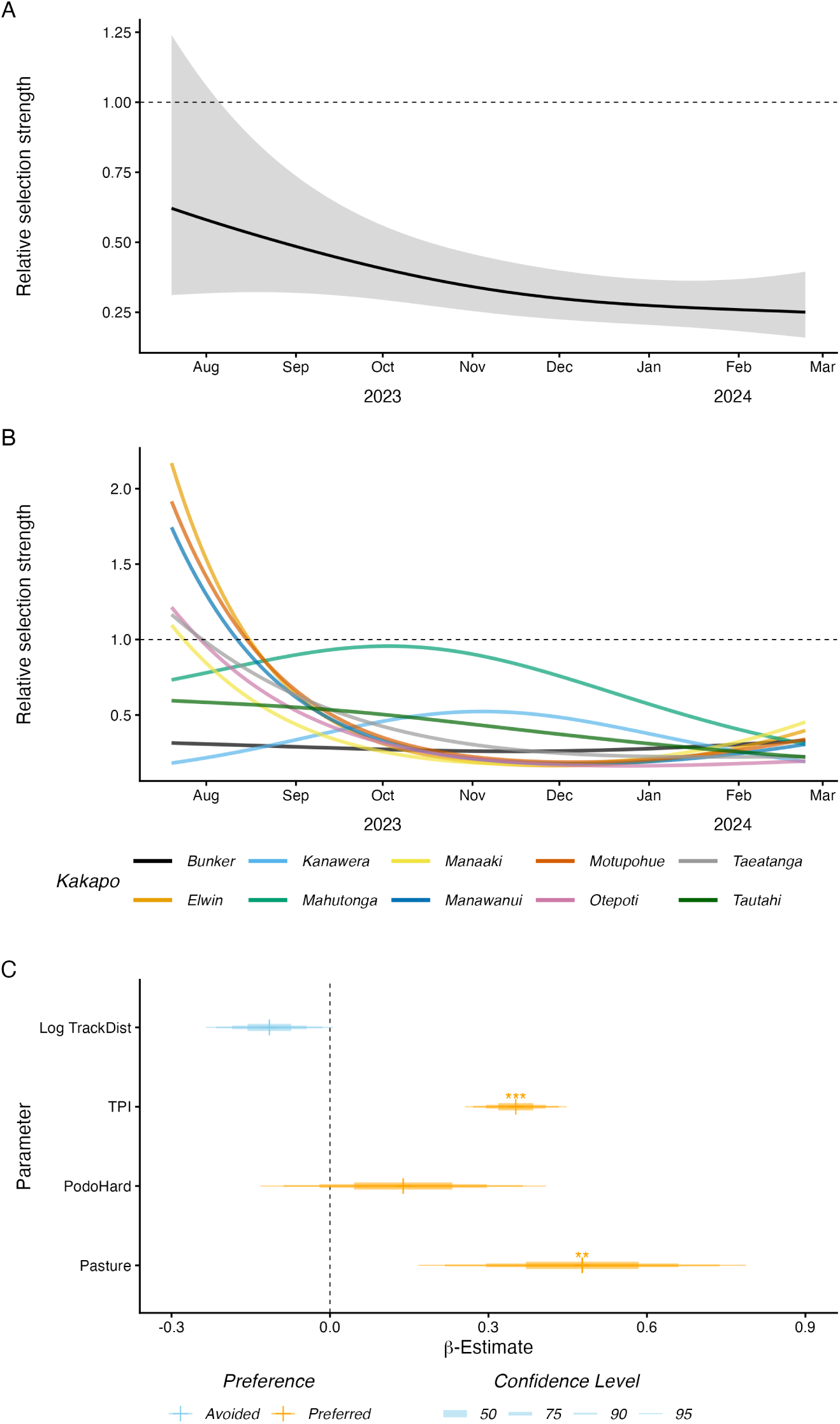
SSF GAM results, with (A) showing the population factor smooth for fence distance with time, (B) individual smooths and (C) the parameter coeflcients. Relative selection strength in (A) and (B) is on the exponential scale (i.e., exp(β_dist_ dist)). For (C), confidence intervals are shown by line thickness in the parameter estimates, with parameters marked with asterisks to represent significance (* = p < 0.05, ** = p < 0.01, *** = p < 0.001).

The resource selection analysis showed that when within 100 m of the fence, kākāpō heavily utilised the small areas of “pasture”, even though this habitat type was relatively scarce within the sanctuary (Figure 4). While only 3% of random locations were within pasture, 40% of observed locations were in this habitat type, showing that pasture was ∼13 times more likely to be selected for than the other vegetation categories (Pearson’s Chi-squared test with Yates’ continuity correction: *p* < 0.001, *χ*^2^ = 2408.6, *df* = 1).

**Figure 4:**
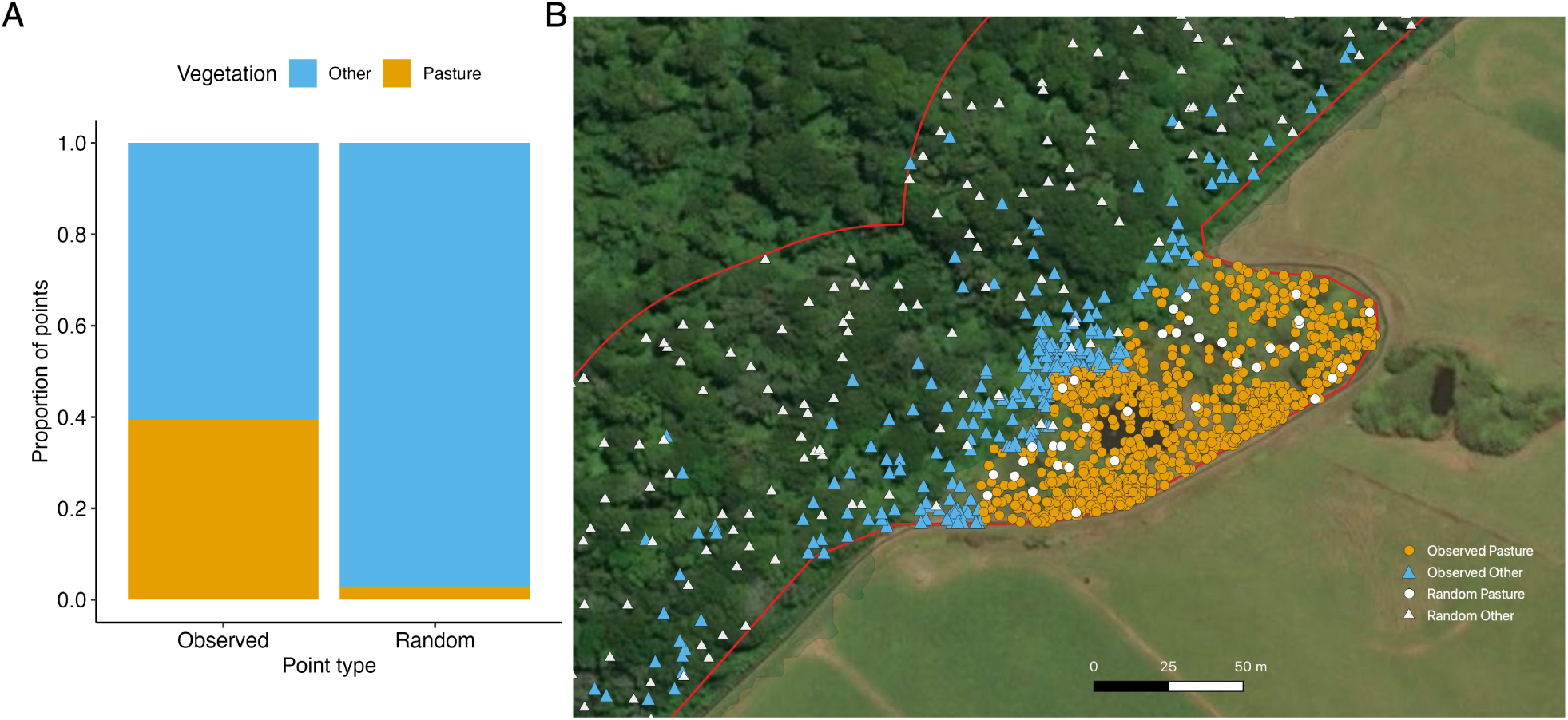
(A) Proportion of observed locations and uniform randomly-sampled points within a 100 m buffer inside the fence that were within pasture compared to other habitat types. (B) a sample map of these points. The random points were uniformly sampled from within the buffer, and were unrelated to the random points paired with observed points used in the SSF analysis. In (B), GPS locations are shown as circles in pasture, and triangles in non-pasture areas. Observed locations are coloured according whether they were in pasture or other areas; random points are all shown as filled white circles (pasture) or triangles (other). Red lines show the fence and 100 m buffer.

## 4. Discussion

The transfer of kākāpō to Maungatautari provided a rare opportunity to use GPS loggers to accurately assess the post-release spatial behaviour of a critically endangered species into a novel environment. That this new site was a fenced sanctuary provided additional information about kākāpō behaviour at this type of protected habitat which has played a crucial role in wildlife conservation in Aotearoa/New Zealand. The behavioural data obtained in this study were important for assessing the success of this kākāpō introduction and informing site selection and future translocations at a time when recovery of the species was limited by habitat availability.

### Main findings

The dynamic space use and dynamic step selection function analyses in this study revealed three unexpected behaviours of the kākāpō at Maungatautari: 1) the space used was larger and movement was more dynamic and exploratory than expected; 2) there was synchronised movement to the fence line; and 3) kākāpō showed strong selection for small patches of exotic habitat, particularly near the fence.

### Large space use

It was expected from previous releases (Joyce, 2009) that the kākāpō would exhibit large movements after release, but that they would settle into smaller home ranges after a short period of one to two months. This did not occur, with high mobility and large areas used by nearly all of the kākāpō over most of the seven month monitoring period. The mean post-release MCP95 of 1,761 ha at Maungatautari in this study is two orders of magnitude larger than estimated ranges of resident kākāpō at other sites, which ranged from 3 – 33 ha in females on Whenua Hou (Whitehead et al., 2011); 15 – 30 ha for males in Fiordland (Best and Powlesland, 1986); and 65 ha per individual on Rakiura / Stewart Island (Powlesland et al., 1995). Joyce (2009) reported larger ranges for males following translocation to Pearl Island, with MCP100 = 84 ha in a non-breeding period and MCP100 = 191 ha in a breeding period, but the ranges found at Maungatautari were still much larger — although the kākāpō on Pearl Island were provided with supplementary food from release which may have reduced their movement, and were on a smaller site with more limited space available for dispersal (an area of 533 ha compared to 3363 ha at Maungatautari). Nevertheless, this spatial behaviour indicates that the kākāpō at Maungatautari remained in a post-release exploratory stage for the duration of the study, and that their behaviour was different to resident kākāpō at other sites. For this reason, the MCP areas at Maungatautari are not representative of settled range sizes at the site, and comparisons with other sites can only be indicative. Furthermore, MCP can be a poor indicator of space use for dispersing animals, and particularly for kākāpō which tend to move quickly between sites in which they settle for a longer period (Trinder, 1998).

### Movement to fence line

The kākāpō at Maungatautari utilised habitat close to the fence more than expected, with marked movement synchronicity between individuals. The occurrence distributions and SSFs showed that most moved towards the fence line simultaneously in November 2023, irrespective of time since their release. The remaining monitored individuals, although few, then mostly moved away from the fence in January 2024. All kākāpō utilised smaller areas during the November to December period, perhaps as a result of reaching the fence barrier and mostly being unable to move beyond it. The synchronicity of movement to the fence line suggests a link to breeding behaviour, since male kākāpō usually move from their winter to summer ranges in spring (Walsh et al., 2005). This downhill movement was the opposite of what was expected, since leks at other kākāpō sites are usually found at high points (Merton et al., 1984). It is unclear why the Maungatautari males did not move uphill, but leks at other sites are often in areas of low scrub or grasslands, which are rare at Maungatautari. Males may have been seeking this or another habitat type or resource not available to them within the sanctuary. However, it should be noted that in the summer following the end of this study (January - February 2025), two of the three remaining males did establish track and bowls and “boom” from them at high points within the sanctuary. During the study, the kākāpō may alternatively have simply been dispersing following translocation and remained at the fence as they tried to move further but were unable to. It may be relevant that the movements towards the fence did not appear random: most of the kākāpō moved to the southern and eastern fence line, with few movements to the north-west (Figure 1). However, there was insufficient information to determine a specific cause of this behaviour.

### Vegetation and feature selection

An alternative explanation of the movement to the fence line is that areas close to the fence provided an attractive habitat for kākāpō. The SSF showed that kākāpō selected for the “pasture” vegetation class more than expected compared to its availability, and that this effect was consistent within 100 m of the fence, where selection for pasture was much greater (∼13 times more) than expected by chance. Although classified as “pasture” in the vegetation map, the vegetation close to the fence in fact contained a variety of plants, including many introduced pasture grasses and shrubs which kākāpō may have not encountered before. An example is blackberry (*Rubus fruticosus*): capture locations and faecal analysis indicate that kākāpō fed on this species both inside and outside the sanctuary. Some small ponds, which do not feature in the vegetation map, also occurred in these areas and may have been an attractant. In addition, the area near the fence line had young fast-growing trees with fresh growth which may have been more accessible and attractive to kākāpō, including both exotic and native species such as tree fuchsia (*Fuchsia excorticata*), wineberry/makomako (*Aristotelia serrata*), mahoe (*Melicytus ramiflorus*), kawakawa (*Piper excelsum*) and willow (*Salix spp.*) (Owen Woodward, Maungatautari, pers. comm.). Some of these native species, such as mahoe and kawakawa, are not present in the southern source range of translocated kākāpō, so their preferences for these species are not known. The possibility that kākāpō selected for the fence area and ridge lines due to the greater diversity of plant species there is supported by foraging habits of remnant kākāpō in Fiordland, where they were attracted to areas of varied seral vegetation associated with recent disturbance (“kākāpō gardens”), alluvial deposits and the treeline (Atkinson and Merton, 2006). However, this “habitat attractant” hypothesis cannot easily provide an explanation for the synchronicity of movement to the fence line in spring — unless kākāpō were attracted to a specific flowering or fruiting event — suggesting that other drivers may have also contributed to this behaviour. Although the use of the pasture patches may have provided an attractant and possibly a supplementary resource, it must also be emphasised that native podocarp/broadleaf forest was the vegetation type most utilised by kākāpō at Maungatautari and it remains the most important known habitat type necessary for their survival.

The SSF also indicated that kākāpō selected for topographic and landscape features, with a weak selection for walking and maintenance tracks and a strong selection for ridge lines, as indicated by topographic position index (TPI). Kākāpō are reported to move along human walking tracks at other locations (Trinder, 1998), presumably because these pathways facilitate movement (Raiter et al., 2018). Ridges may have been used for easier travel (compared to thickly-vegetated gullies), but also because the males might have been investigating potential display sites at higher elevation. However, the selection for ridge lines may have been due to GPS reception bias, as explained below. Another potential confounding factor is that some tracks follow ridge lines, but this is unlikely to be able to fully explain the strength of the TPI effect since there was only a weak correlation between track distance and TPI (Spearman rank correlation *r_S_* = −0.27). The ridge lines may have also offered better feeding opportunities for kākāpō, where the lower and more open canopy would be more accessible to kākāpō and provide greater variation of palatable species, including subcanopy species such as toropapa (*Alseuosmia macrophylla*), and grasses such as gahnia (*Gahnia spp.*) which are less common as lower elevations (Owen Woodward, Maungatautari, pers. comm.).

### Limitations

The conclusions reached in this study should be considered alongside the constraints and limitations of the data. The study only assessed short-term movement patterns in the seven month period immediately after translocation, during which there was large variation in monitoring success among individuals (Figure S2). Longer periods of monitoring would be necessary to assess kākāpō behaviour as they settle into the residency stage, particularly during the spring and early summer period (November – December) when the individuals showed simultaneous movement that was potentially associated with breeding behaviour.

Estimated habitat selection may also have been affected by non-uniform GPS reception across the site. GPS fix success is likely to have been lower in gullies and in thickly forested areas, where the tags would have had a lower area of sky visible and fewer satellites available. This may have contributed to the apparent kākāpō selection for ridges and pasture or scrub. However, this bias is unlikely to have been fully responsible for the large TPI effect seen, especially since topographic obstruction has been shown to have a lower effect on GPS accuracy than canopy closure (which may only become prominent at > 90% canopy closure; Forrest et al., 2022). The magnitude of pasture selection also suggests that this bias is unlikely to account for the entire effect.

Another consideration is that the Kriging approach for estimating occurrence distributions (ODs) did not consider the fence as a barrier, which is possible with some range estimators (e.g. Hollins et al. 2024). Although truncating the ODs at the fence may have produced more realistic space use estimates, we do not expect that it would have changed the conclusions from our results. Our results are more likely conservative, as truncating space use estimates at the fence line would likely have strengthened the trend of a contraction of space use that was closer to the fence.

### Conservation implications and future study

The results from this study suggest that kākāpō at Maungatautari were unable to settle at the site in the seven months following release. This could indicate that the kākāpō may have been unable to find suitable habitat to meet their dietary or behavioural needs, or that they needed more time to adapt to their new habitat. We suggest that there is more support for the latter explanation, as the largely intact native forest at Maungatautari should be sufficient for kākāpō to maintain non-breeding weights. This was consistent with the kākāpō being in good health during the study period, with minimal post-transfer weight loss recorded during regular health checks (which occurred 25 times in total, excluding transfers). Further, the selection for pasture and exotic vegetation at the fence line also supports this hypothesis, as does the potential attractant to ridge lines due to novel and more diverse and accessible vegetation. Kākāpō from the same source population had previously been transferred to another novel habitat on Hauturu-o-Toi/Little Barrier Island, which is further north and has a greater contrast in vegetation and climate compared to Maungatautari. The kākāpō appeared to settle well there, but those individuals were only infrequently monitored and successful breeding tended to only be possible when supported by supplementary feeding (Stone, 2013; Stone et al., 2017). Hauturu-o-Toi also did not have the range of exotic introduced vegetation available at Maungatautari. The possibility that kākāpō released at sites with different habitat from their source site may need longer to acclimatise should be considered during translocation planning and monitoring.

It is also possible that the behavioural requirements of kākāpō at Maungatautari in relation to breeding were not met, and that the males were moving to find suitable display sites or other kākāpō (either males or females). The synchronicity of movement to the fence at the start of the usual breeding period in late spring supports a reproductive basis for their unexpected spatial behaviour. The low density of kākāpō at Maungatautari may have exacerbated this movement, with densities of up to one kākāpō per 300 ha compared to as little as one per 15 ha at their source sites. Animal dispersal is affected by conspecific density in a wide range of species, although responses are dependent on a range of factors such as life history stage and even individual personality (Jreidini and Green, 2024) — which may partially explain the individual variation in movement in this study. Although it is not possible to definitively determine from this study if breeding requirements influenced spatial behaviour, the synchronisation of movements in spring strongly suggest that timing may be an important factor to consider in translocations, and that future transfers should be conducted in a manner to help investigate and account for this behaviour.

Alternatively, the large movements exhibited by kākāpō in this study may be normal following translocation, and either previous post-transfer monitoring using radio telemetry has not been of sufficient temporal and spatial resolution to reveal this behaviour, or there was less space for dispersal at other sites. Joyce (2009) provided the most comprehensive post-translocation monitoring of kākāpō to date, with 4424 fixes of 26 individuals (13 males and 13 females), albeit with lower resolution radio telemetry. This study found that individuals did tend to exhibit greater movement during a dispersal stage, which lasted a mean of 14 days (range = 0 – 38 days) for males and 45 days (range = 7 – 185 days) for females, although the female mean reduced to 33 days (range = 7 – 58 days) if a single female was excluded. Following release in April 1998, all 13 males had established display sites by the breeding period in late 1998. Even allowing for the lower spatial and temporal resolution of the kākāpō locations, these results indicate that kākāpō at Maungatautari would have been expected to settle into stable ranges during the ∼seven month period of the current study. However, the translocation studied by Joyce (2009) was to a much smaller island site (Pearl Island, with an area of 533 ha compared to 3363 ha at Maungatautari) which had very similar climate and vegetation to the nearby source island (Whenua Hou/Codfish Island), and included females as well as males just prior to a breeding season. This suggests that habitat variation, site size, access to conspecifics, timing relative to breeding periods or a combination of all of these were more likely to have led to the much unexpectedly long dispersal period at Maungatautari.

## 5. Conclusions

The translocation of kākāpō to Maungatautari in 2023 provided a natural experiment to examine the post-translocation habitat use of a critically endangered species in an environment novel to the contemporary relict population. High-resolution GPS monitoring was used to improve the limited understanding of kākāpō spatial behaviour across different environments. Dynamic space use and step selection function analyses revealed several unexpected behaviours, including extensive movements during a prolonged exploratory dispersal stage, synchronised movements to the perimeter fence, and unexpected selection for exotic vegetation. These findings suggest that translocated kākāpō may respond to novel environments in ways that are difficult to predict from observations of established populations.

The reasons for these large and synchronised movements are unclear, but are likely to be related to breeding behaviour, the distribution of conspecifics, or the availability of familiar habitat features. Future work should focus on continued monitoring of the Maungatautari population to determine whether these behaviours persist as individuals become established, and on comparisons with other recent translocations to evaluate the generality of these patterns. This research will help inform the planning, implementation and evaluation of future translocations, which are a high priority for kākāpō recovery. More broadly, these results highlight the value of long-term, high-resolution spatial monitoring for the conservation management of threatened species.

## Declaration of generative AI and AI-assisted technologies

The authors used the generative artificial intelligence tool Microsoft365 CoPilot (powered by the GPT-5 model) for minor coding assistance during the preparation of this manuscript. The results were manually verified, and the authors take full responsibility for the content of the published article.

## Acknowledgments

We are grateful to the staff of Sanctuary Mountain Maungatautari, the Kākāpō Recovery Team (KRT) and Auckland Zoo for the fieldwork and associated support which made this study possible, particularly Dan Howie, Craig Montgomerie, Janelle Ward, Bodie Taylor, Helen Hughes, Tori Budd and Caitlin Stoyel-Wiggens from Maungatautari; Daniella Whitaker, Petrus Hedman and Tommy McKerras from KRT, Rose Graham from NZDOC; and Jasmine Rabaud and Mikaylie Wilson from Auckland Zoo. Eeva Kumpula assisted with data cleaning, Nick Perham with GIS data manipulations, Luke Lloyd Jones with SSF-GAM modelling and Owen Woodward provided knowledge of vegetation at Maungatautari necessary to infer relevance of kākāpō behaviour. Geoff Churchill contributed fence and track GIS data, and GIS In Conservation provided other GIS shapefiles.

The authors are grateful to indigenous Ngāi Tahu whānui for supporting and enabling the test of kākāpō habitat outside of the Ngāi Tahu takiwā (region). A central component of this transfer was an iwi whanaungatanga (relationship) agreement to support the handing of kākāpō from Ngāi Tahu to the mana whenua (customary landholders): Ngāti Korokī Kahukura, Ngāti Hauā, Ngāti Raukawa and Te iwi o Tainui. This ensured that kawa, tikanga (customs), kaitiakitanga (guardianship) and manaakitanga (hospitality) were used to provide assurances to both Ngāi Tahu and Tainui that the mauri (life force) of kākāpō remained safe on the maunga (mountain).

The authors appreciate the contribution of those who worked for many years to achieve the translocation of kākāpō to Maungatautari, and in particular to Chris Smuts-Kennedy and John Innes.

## Funding

this work was supported by the Kākāpō Recovery Programme (supported by Meridian Energy Ltd and public donations through the Mauri Ora Kākāpō Trust), Sanctuary Mountain Maungatautari, the World Parrot Trust, the New Zealand Parrot Trust and On The Edge Conservation, UK. SW Forrest helped to prepare the manuscript for publication while affiliated with Queensland University of Technology and CSIRO, and acknowledges support by an Australian Government Research Training Program Doctoral Scholarship and a CSIRO top-up scholarship. Open access publishing facilitated by University of Tasmania, as part of the Wiley - University of Tasmania agreement via the Council of Australian University Librarians.

## Author Contributions

- Conceptualisation: AD, DE, DV, SF, TD.
- Data curation: AD, DE, DV, KRT, DH, CM, JW.
- Formal analysis: SF, AD, NK.
- Visualisation: AD, SF, NK.
- Writing - original draft: AD, SF.
- Writing - review and editing: AD, SF, NK, DE, DV, TD, DH, JW, CM.
- Funding: DV, AD, DE, TD.

## Collaborative Author Groups

The Kākāpō Recovery Team staff contributing to this research were: Andrew Digby, Daryl

Eason, Andie Gentle, Petrus Hedman, Bryony Hitchcock, Scott Latimer, Sarah Little, Sarah Mank-telow, Tommy McKerras, Jake Osborne, Brodie Philp, Louise Porter, Alyssa Salton, Polina Stucke, Theo Thompson, Lydia Uddstrom, Deidre Vercoe, Alex Webster, Daniella Whitaker, Maddy Whittaker.

## Supporting Information

### A. Data collection

#### A.1. GPS schedule

**Figure S1:**
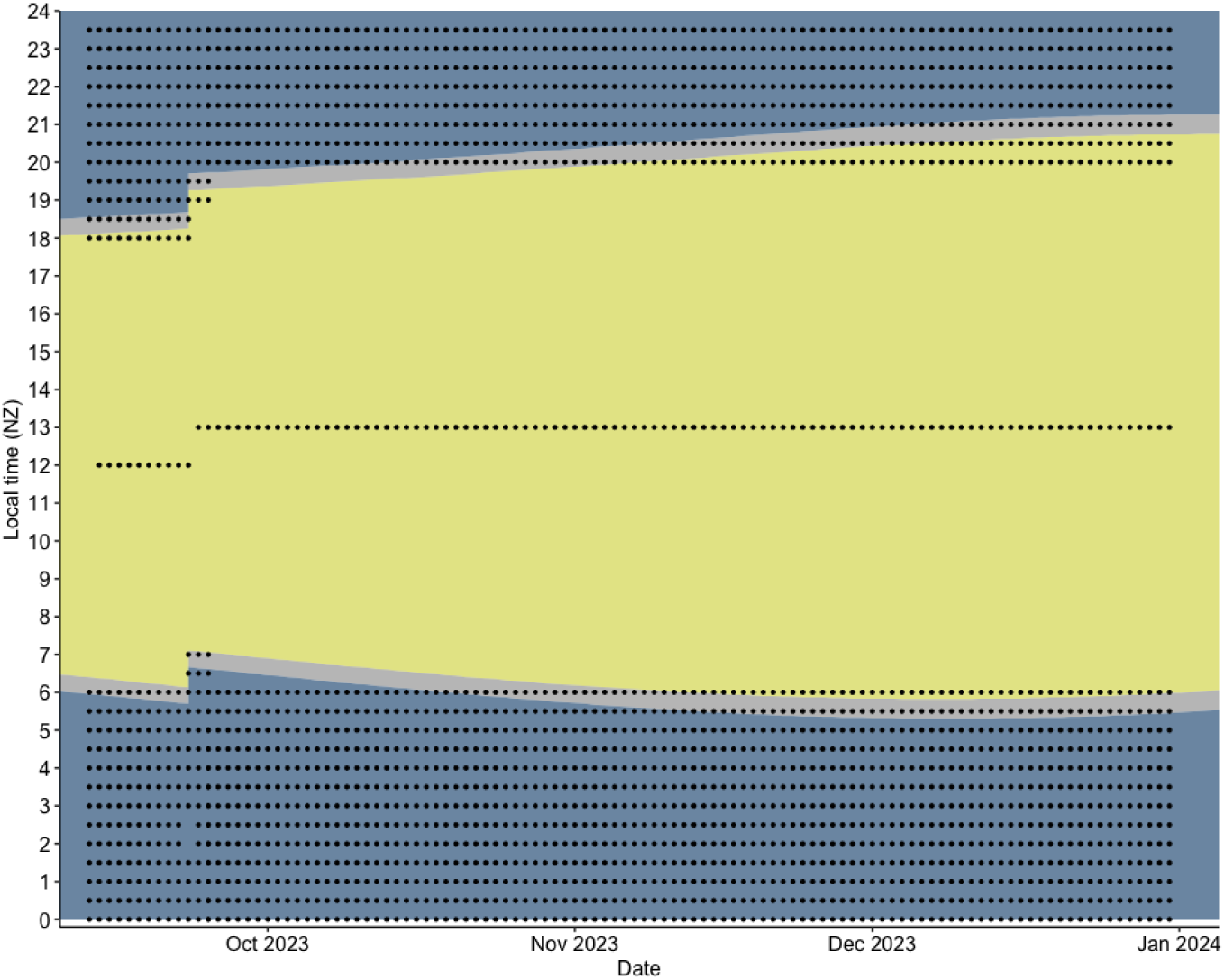
Fix schedule for the Lotek PinPoint 240 GPS tags deployed from September 2023 to Jan 2024. Each black dot represents a programmed fix attempt. Daylight is shown as yellow shaded area, night as blue, and civil twilight as grey.

**Figure S2:**
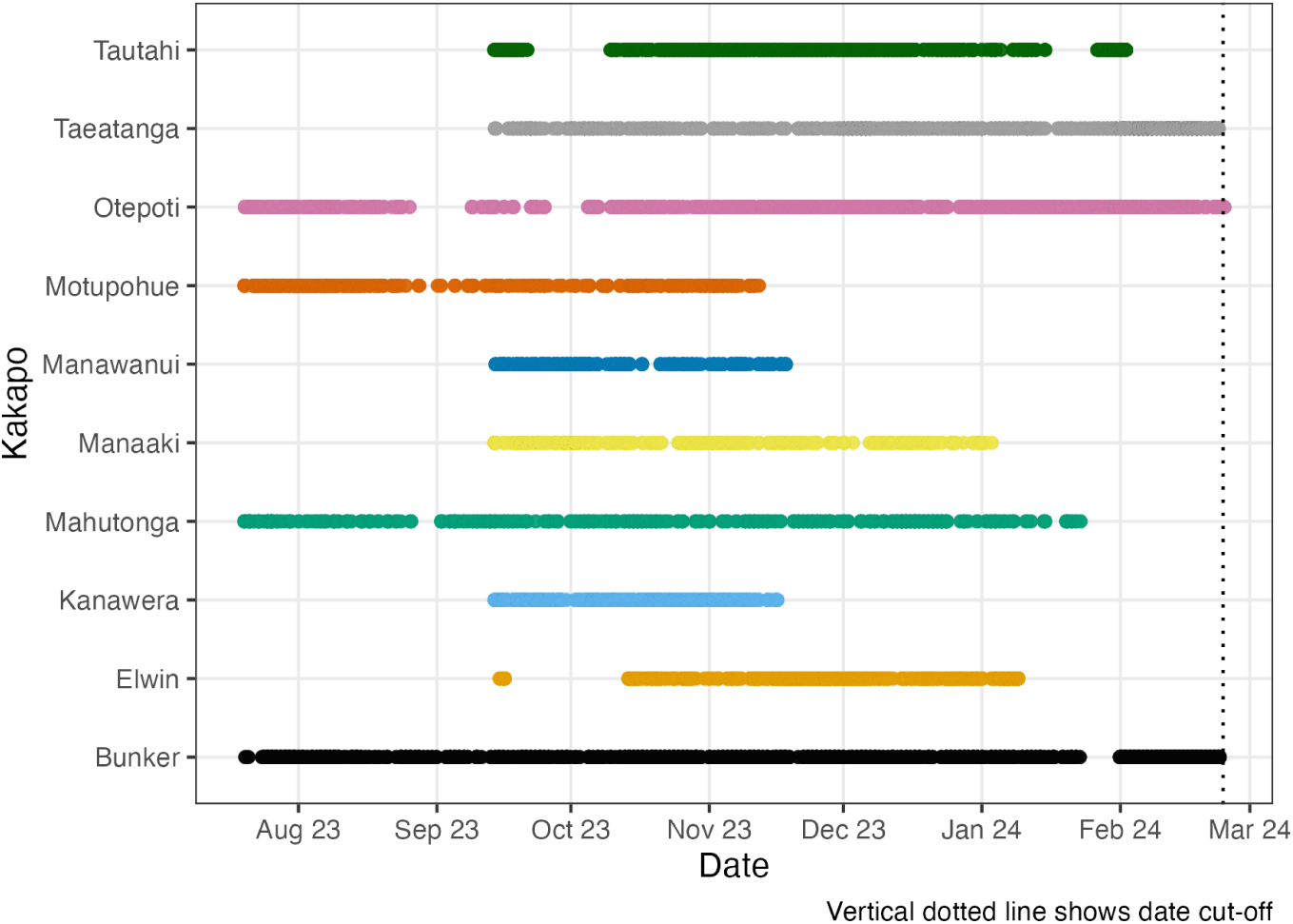
Timeline of GPS data collected for each kākāpō. The vertical dotted line denotes the last date of 24th Feb 2024 used in the analyses.

**Figure S3:**
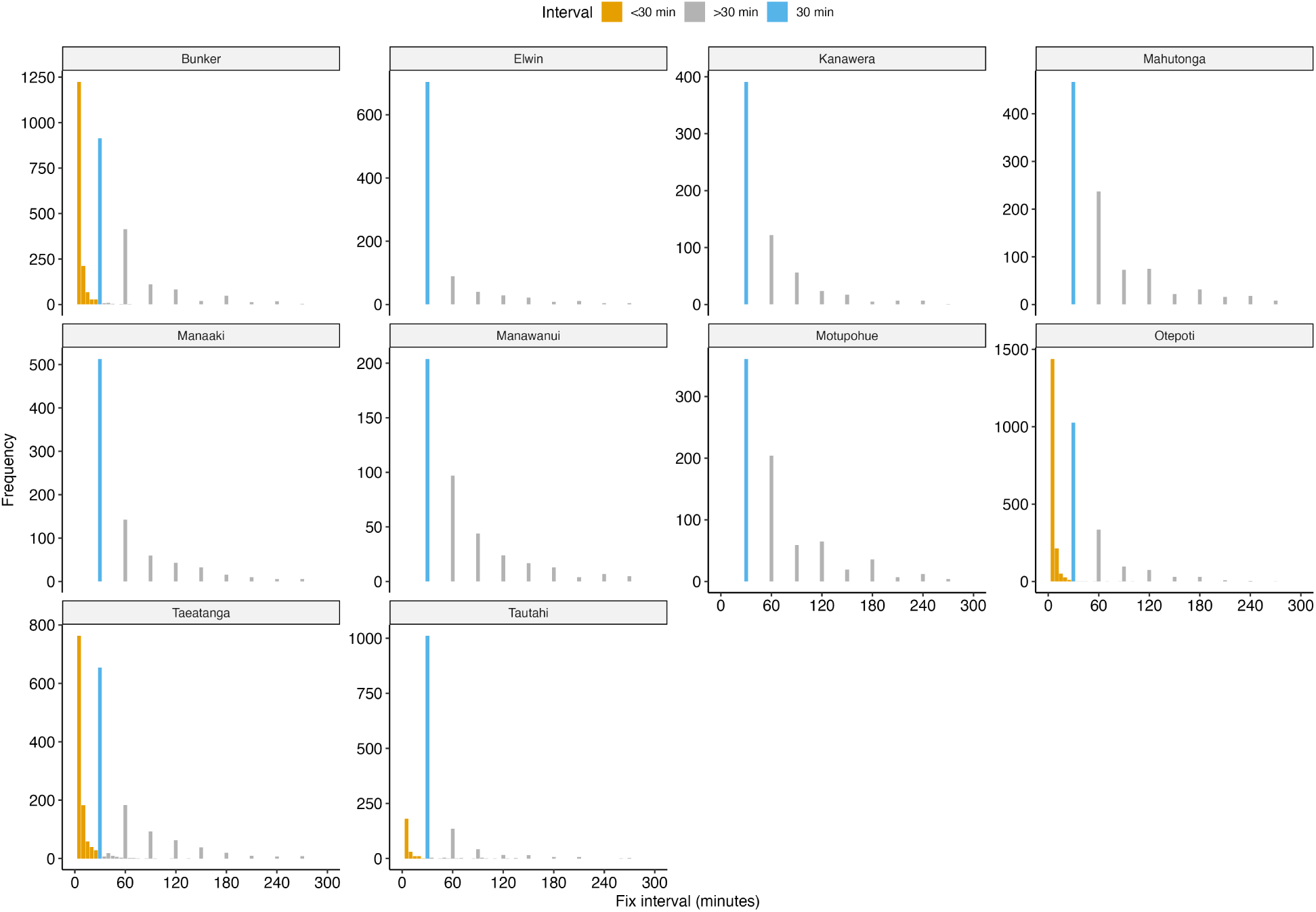
A histogram of the interval between GPS fixes for each individual kākāpō. Blue denotes the standard 30 minute fix interval used in most cases; gold indicates the fixes from the five minute schedule used for four individuals. Fixes with all interval values were used in the SSF, but those with fix intervals less than 30 minutes were resampled to 30 minute intervals for the dynamic space use analysis.

### B. Data preparation

#### B.1. Speed filtering

Location errors were removed using speed filtering. The maximum linear speed between consecutive fixes for each individual kākāpō was derived using the function vmax in package SDLFilter (Shimada et al., 2012) in R (R Core Team, 2025). We used a maximum likelihood estimate to fit a Gamma distribution to the speeds between all locations derived from at least 5 satellites, and set the linear speed threshold per individual kākāpō as the 99th percentile value of this distribution. A speed threshold was set separately for each kākāpō over the entire monitoring period to account for individual variation in behaviour. Locations were then removed from the dataset if the linear speed before OR after a fix was greater than the per-individual threshold, regardless of the number of satellites or HDOP (Forrest et al., 2024).

#### B.2. GPS data

**Table S1:** The GPS dataset for the ten male kākāpō at Maungatautari, combined over multiple deployments.

| Kākāpō | Age<sup>a</sup> | Attempts<sup>b</sup> | Fixes<sup>c</sup> | Vmax<sup>d</sup> | FSR (%)<sup>e</sup> | Days<sup>f</sup> | Start<sup>g</sup> | End | DSU<sup>h</sup> | SSF<sup>i</sup> | MCP95<sup>j</sup> |
| --- | --- | --- | --- | --- | --- | --- | --- | --- | --- | --- | --- |
| Bunker | J | 6425 | 3468 | 1.04 | 54 | 209 | 19/07/23 | 23/02/24 | 2188 | 2388 | 803 |
| Elwin | A | 2056 | 1052 | 0.64 | 51 | 91 | 14/09/23 | 10/01/24 | 1031 | 592 | 712 |
| Kanawera | A | 1457 | 708 | 0.66 | 49 | 63 | 13/09/23 | 16/11/23 | 707 | 513 | 2319 |
| Mahutonga | J | 3895 | 1237 | 0.30 | 32 | 187 | 19/07/23 | 24/01/24 | 1210 | 709 | 766 |
| Manaaki | J | 2531 | 1018 | 0.62 | 40 | 112 | 13/09/23 | 04/01/24 | 952 | 476 | 2916 |
| Manawanui | J | 1504 | 506 | 0.53 | 34 | 64 | 14/09/23 | 18/11/23 | 500 | 263 | 1756 |
| Motupohue | J | 2347 | 924 | 0.85 | 39 | 115 | 19/07/23 | 13/11/23 | 904 | 588 | 1762 |
| Otepoti | J | 6835 | 3765 | 0.84 | 55 | 218 | 19/07/23 | 24/02/24 | 2192 | 2544 | 2450 |
| Taeatanga | A | 5592 | 2393 | 1.38 | 43 | 161 | 13/09/23 | 22/02/24 | 1532 | 1788 | 2503 |
| Tautahi | J | 3608 | 1943 | 0.61 | 54 | 132 | 13/09/23 | 02/02/24 | 1409 | 1057 | 1818 |
| **Total/mean** |  | 36250 | 17014 | 0.747 | 45.1 | 1352 |  |  | 12625 | 10918 | 1761 |

<sup>a</sup> Age class: A = adults (at least 5 years old); J = juvenile (under 5 years old). <sup>b</sup> The number of fix attempts. <sup>c</sup> The number of successful fixes from those attempted after duplicates were removed and Vmax filtering applied. <sup>d</sup> Maximum speed per individual used for speed filtering, in km/h <sup>e</sup> The percentage fix success rate (FSR) after Vmax filtering, defined as $100 \times \text{Fixes Vmax} / \text{Fixes attempted}$ . <sup>f</sup> The combined length of the deployments between these dates. <sup>g</sup> Start and end date show the first and last dates on which fixes were successful, after filtering. <sup>h</sup> The number of locations used in the dynamic space use analysis. <sup>i</sup> The number of locations used in the step selection function analysis. <sup>j</sup> MCP95 areas calculated over all time periods from the step selection function dataset.

#### B.3. Vegetation classes

**Table S2:**
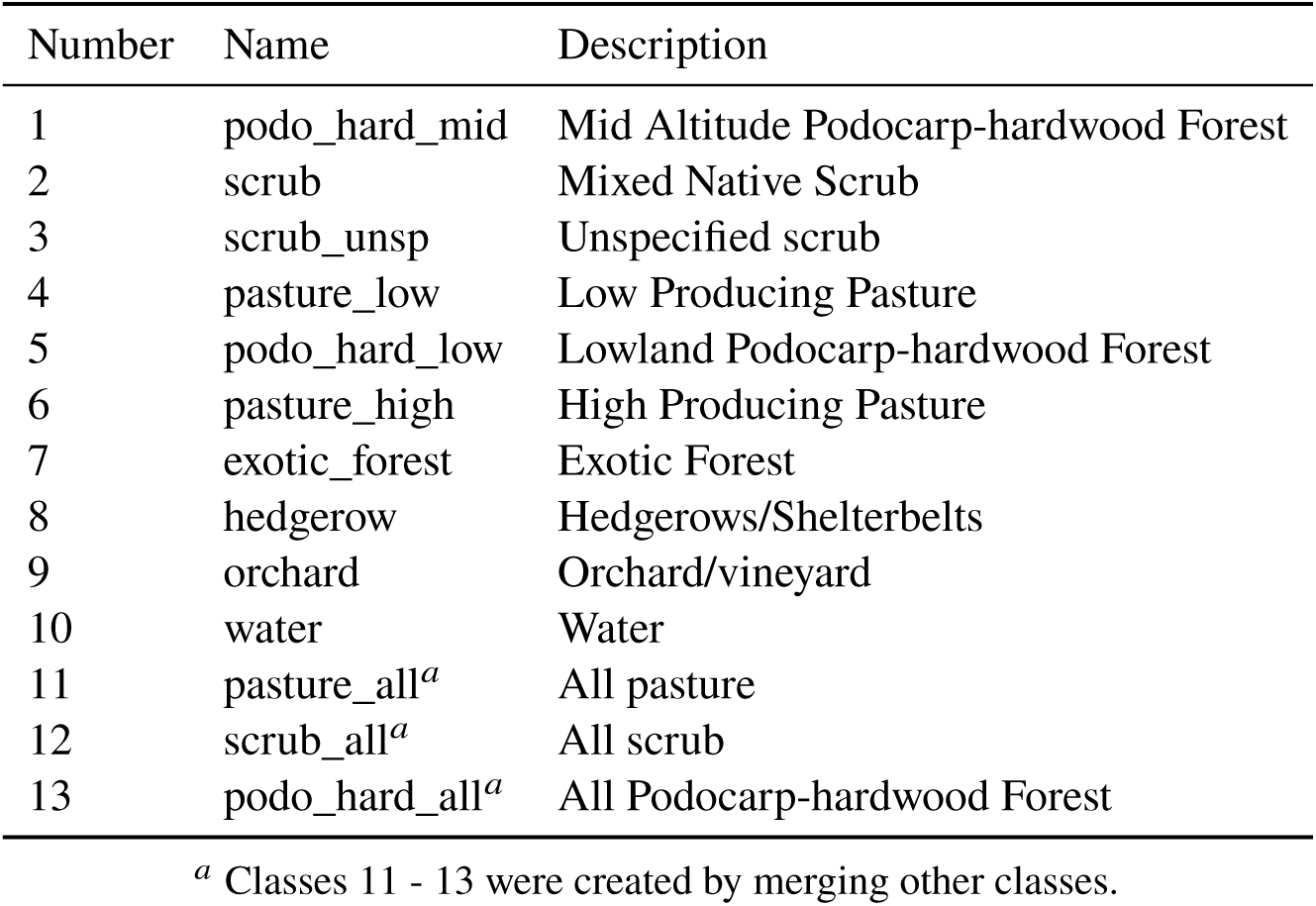
Vegetation classes in the full vegetation map which were considered for the step selection function analysis.

#### B.4. Fence reflection

Figure S4 shows the distribution of locations when the kākāpō were known to be inside the fence from independent radio telemetry observations. The distribution is centred just inside the fence, with a symmetric and long-tailed distribution inside and outside of the fence. This suggests that for these locations, the kākāpō were near the fence but that GPS error caused the locations to be placed inside and outside of the fence. This is supported by the distribution divided by number of satellites for each fix: those with higher numbers of satellites, assumed to be more accurate, are closer to the fence (Figure S5). To prevent removing a large number of fixes that were just outside of the fence, we reflected the distance of locations that were 1 – 200 m outside the fence to the same distance inside the fence, maintaining the GPS error distribution.

**Figure S4:**
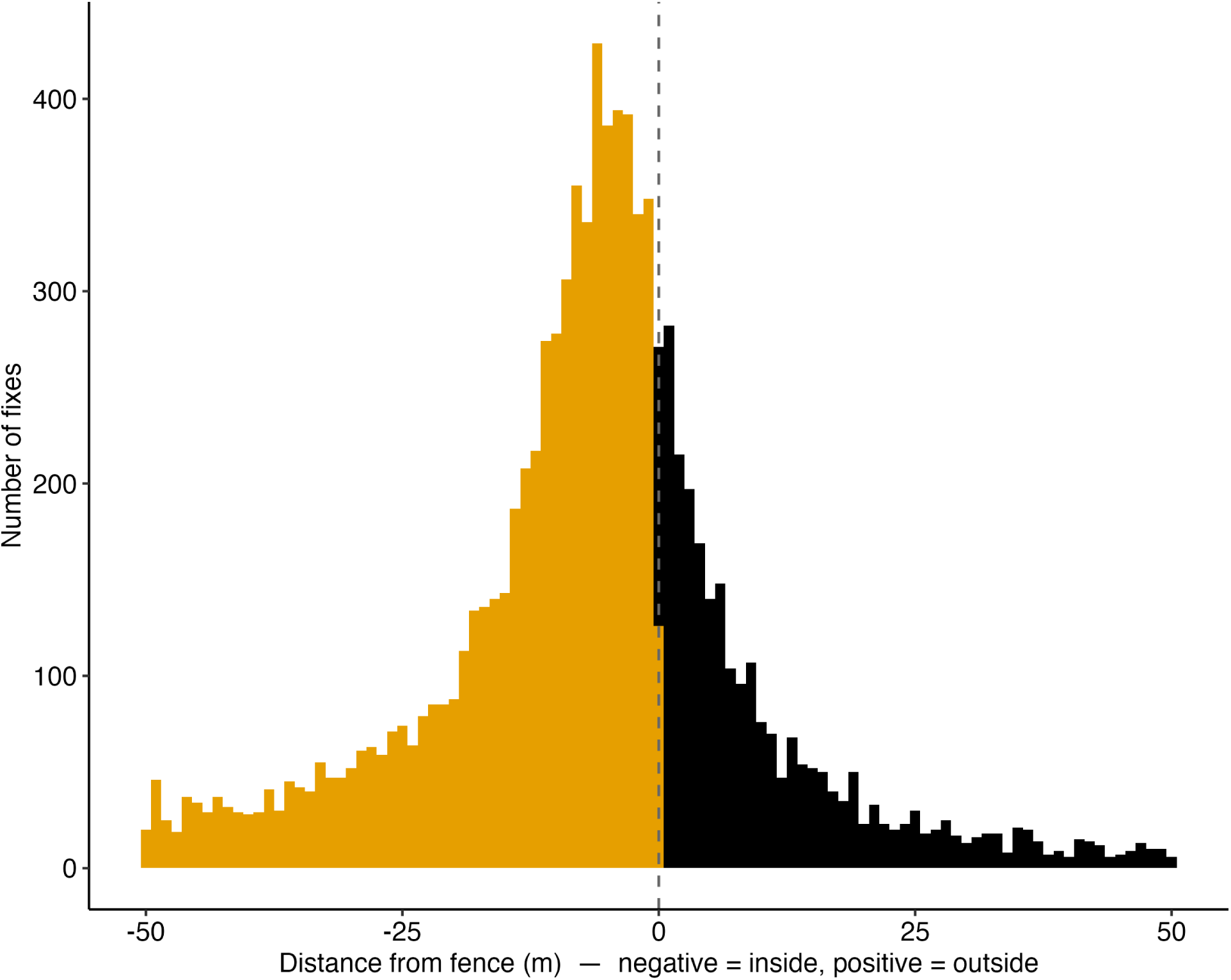
The distribution of location distances inside and outside the fence, for when the kākāpō were known to be inside the fence from independent radio telemetry observations.

**Figure S5:**
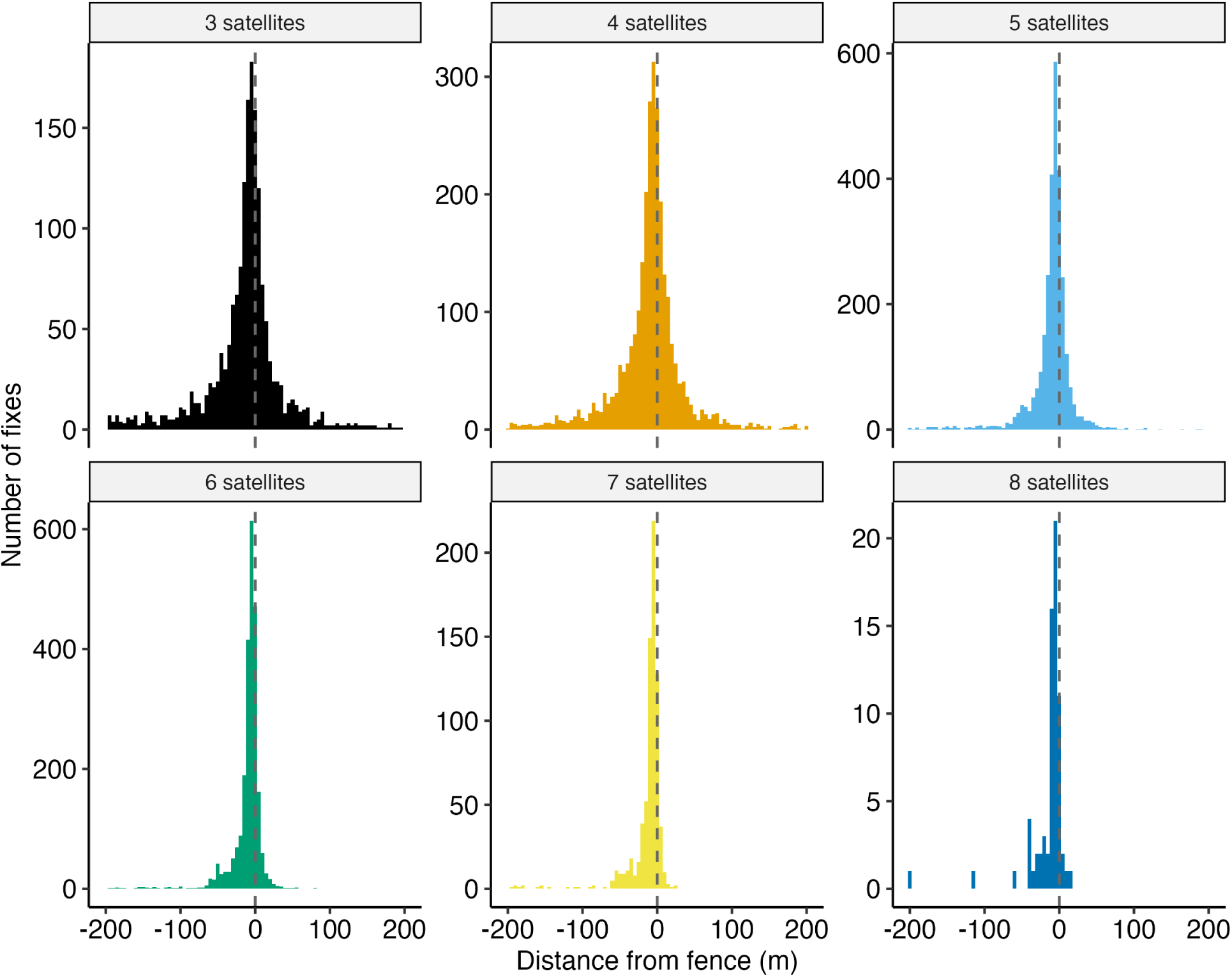
The distribution of location distances inside and outside the fence for increasing numbers of satellites. For locations with higher numbers of satellites, which we assume are more accurate, then we see that many of those locations (i.e. six to eight satellites) are very close to the fence.

### C. Step selection functions

#### Observed and random step distributions

**Figure S6:**
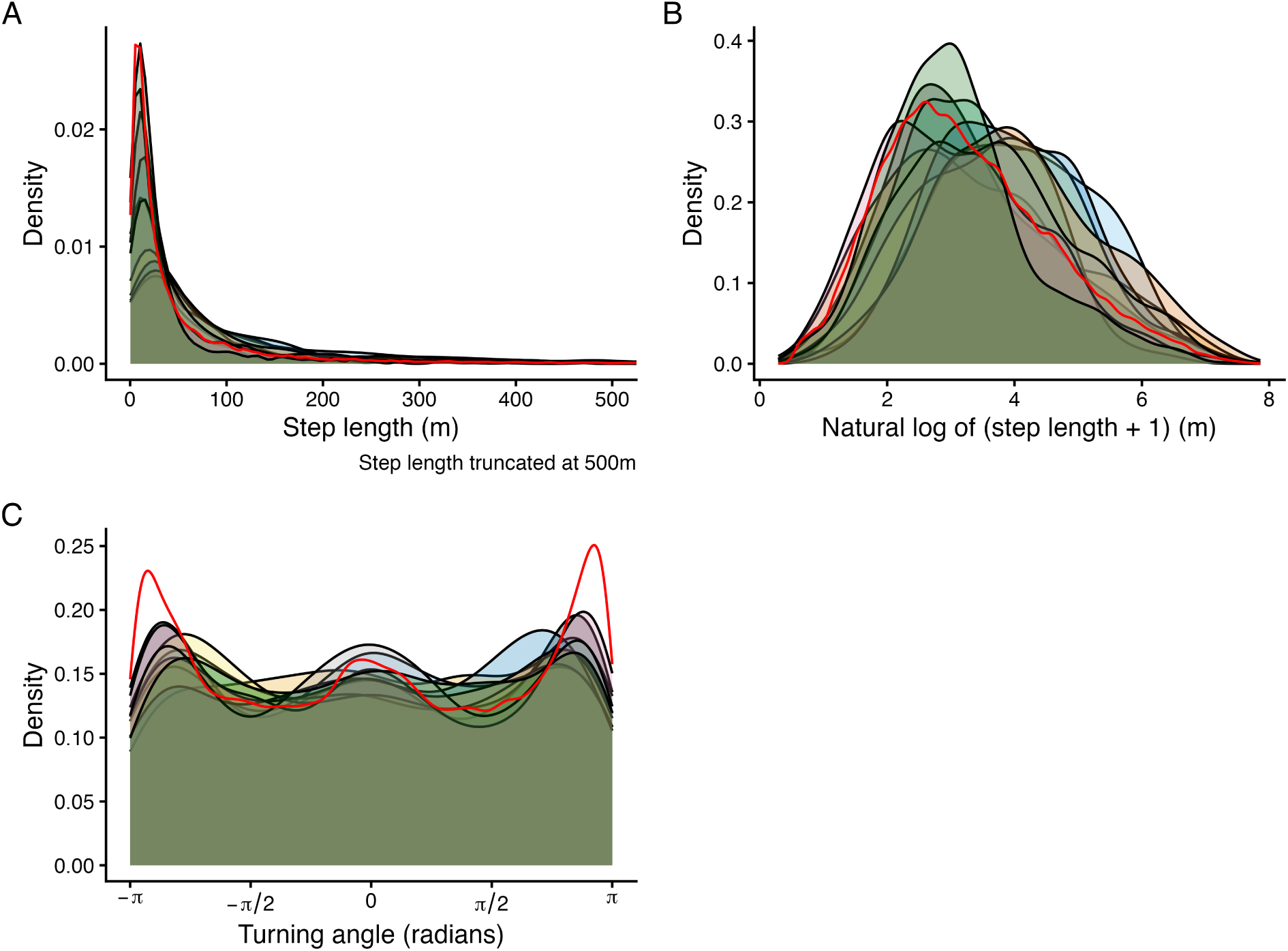
The distribution of step lengths (A), log of step lengths (B) and turning angles (C) for random steps (red line) and observed steps (shaded areas, with a different colour for each kākāpō). There was little evidence of multimodality when log-transforming the step lengths. Peaks of turning angles near 0 and at −π/π indicated that movement was not highly directed.

#### C.1. Topographic Position Index (TPI) radius selection

The radius for TPI was determined by setting a range of values from 100 m to 2500 m (100 m, 300 m, 500 m, 1000 m and 2500 m) and including each TPI as a covariate along with fence distance, track distance and vegetation, and then selecting the best-performing SSF using the corrected Akaike Information Criterion (AICc) calculated by function aictab in package AICcmodavg (Mazerolle, 2023). This analysis showed that TPI with a radius of 300 m best explained the data (AICc weight= 0.965), so this was used as a covariate in the final SSF (Figure 1D).

**Figure S7:**
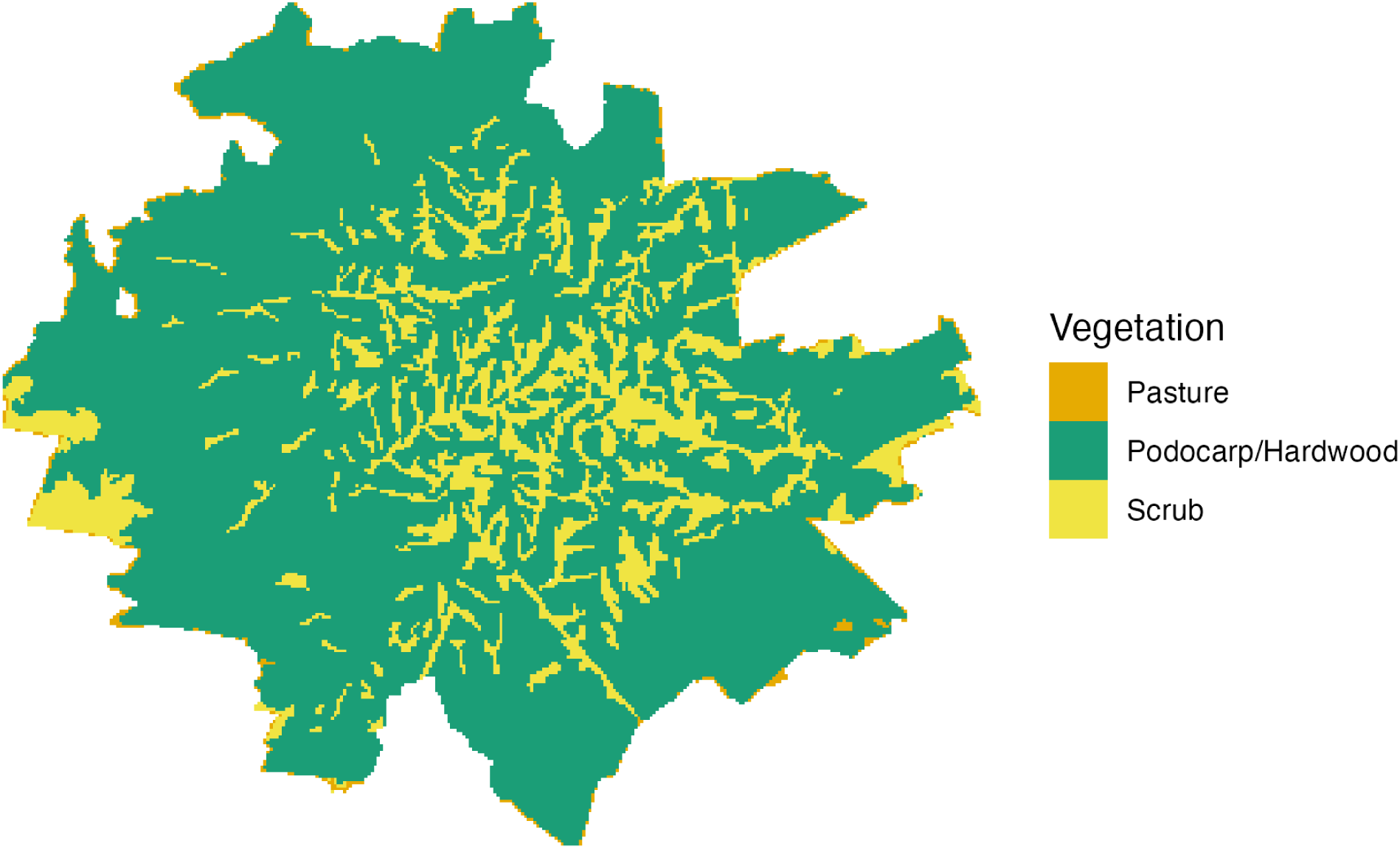
Map of the vegetation classes considered for the step selection function models within the fence which were used in the SSF models: pasture, podocarp forest and scrub. Note that within the sanctuary there are only small patches of pasture which are mostly adjacent to the fence line.

**Figure S8:**
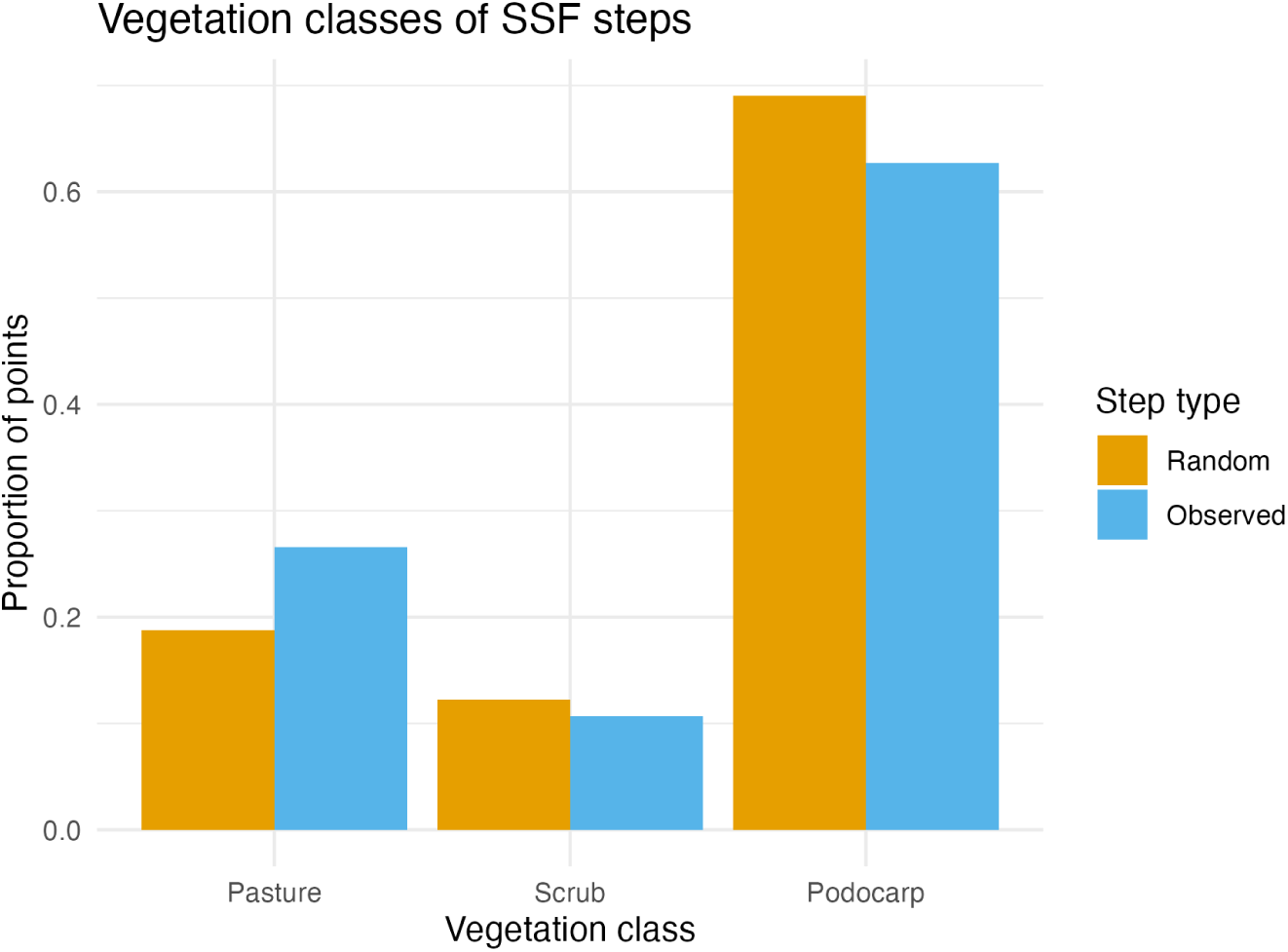
The proportion in each vegetation class of the observed and random steps used in the SSF. The proportion of both random and observed steps sum to one, but the frequency of random steps was higher — there were 20 random steps to every observed step. The native vegetation class (podocarp) was the most highly used vegetation class, partly owing to its high prevalence throughout Maungatautari. Although pasture only constituted a very minor proportion of the overall landscape, more than 20% of the observed kākāpō locations fell within it.

#### C.2. SSF validation

**Figure S9:**
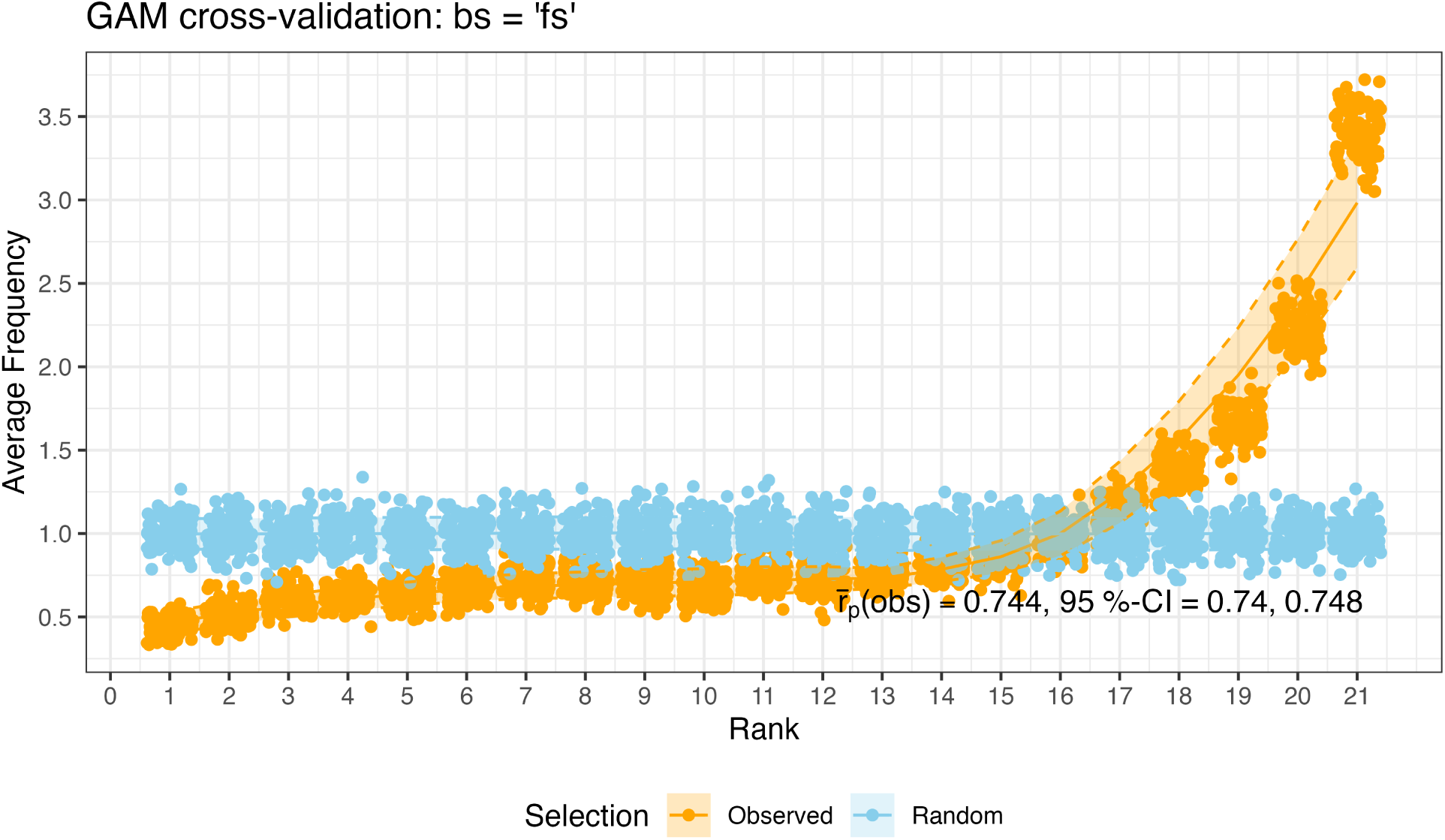
Cross-validation results for the final model that used a factor smooth term to account for individual-level random effects. We re-ran the model 100 times, where 80% of the data was used for model fitting, and 20% was used for testing (selected randomly, preserving strata). For each replicate, step selection weights for the observed and random steps were calculated using the estimated parameters of the full model using the predict.gam function from the mgcv package (Wood, 2011). The weights of the 21 steps (1 observed and 20 random) were ranked, where a rank of 1 is the lowest weight (least likely step), and a rank of 21 has the highest weight (most likely step). The y-axis shows the average frequency of the ranks for the observed steps (orange) and a step that was selected randomly (blue), with one point per rank and cross-validation replicate (n = 100). In this case, the observed step was the highest ranked step at an average frequency of 3.4, meaning that the observed step had the highest probability weight 3.4 times more than would be expected by chance. The Pearson correlation coeflcient (r_p_) was calculated for the rankings (1–21) and their associated frequency, and the average r_p_ ± 95% confidence intervals is shown for the observed steps. Values of r_p_can range from –1 (negative linear correlation) to 1 (positive linear correlation), with 0 indicating a model performs as well as random chance.

**Figure S10:**
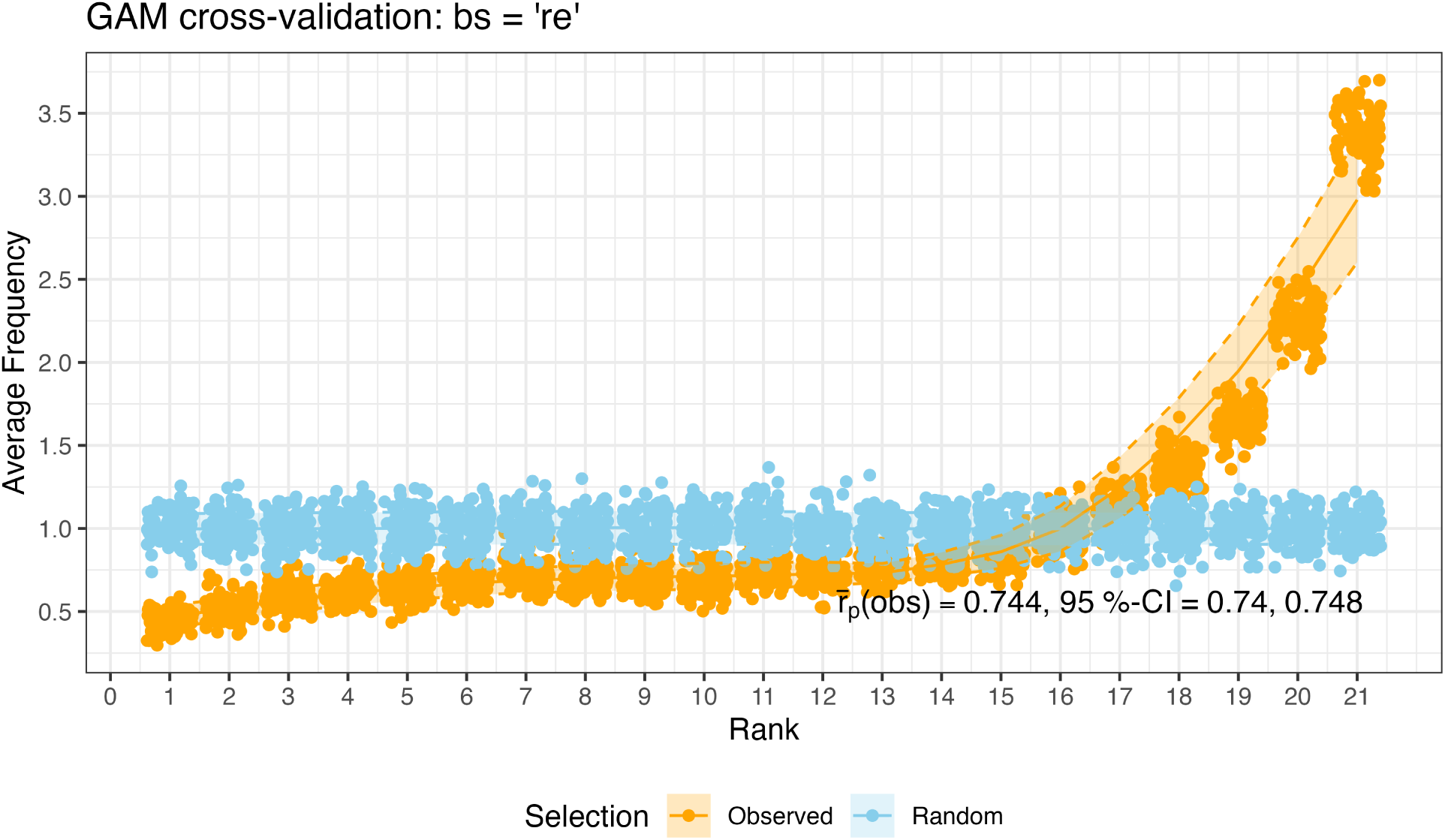
Cross-validation results for the alternative model that used a random slope to account for individual-level random effects. The ‘Random’ model is a null model where the rank was selected randomly from the observed and proposed steps.

**Figure S11:**
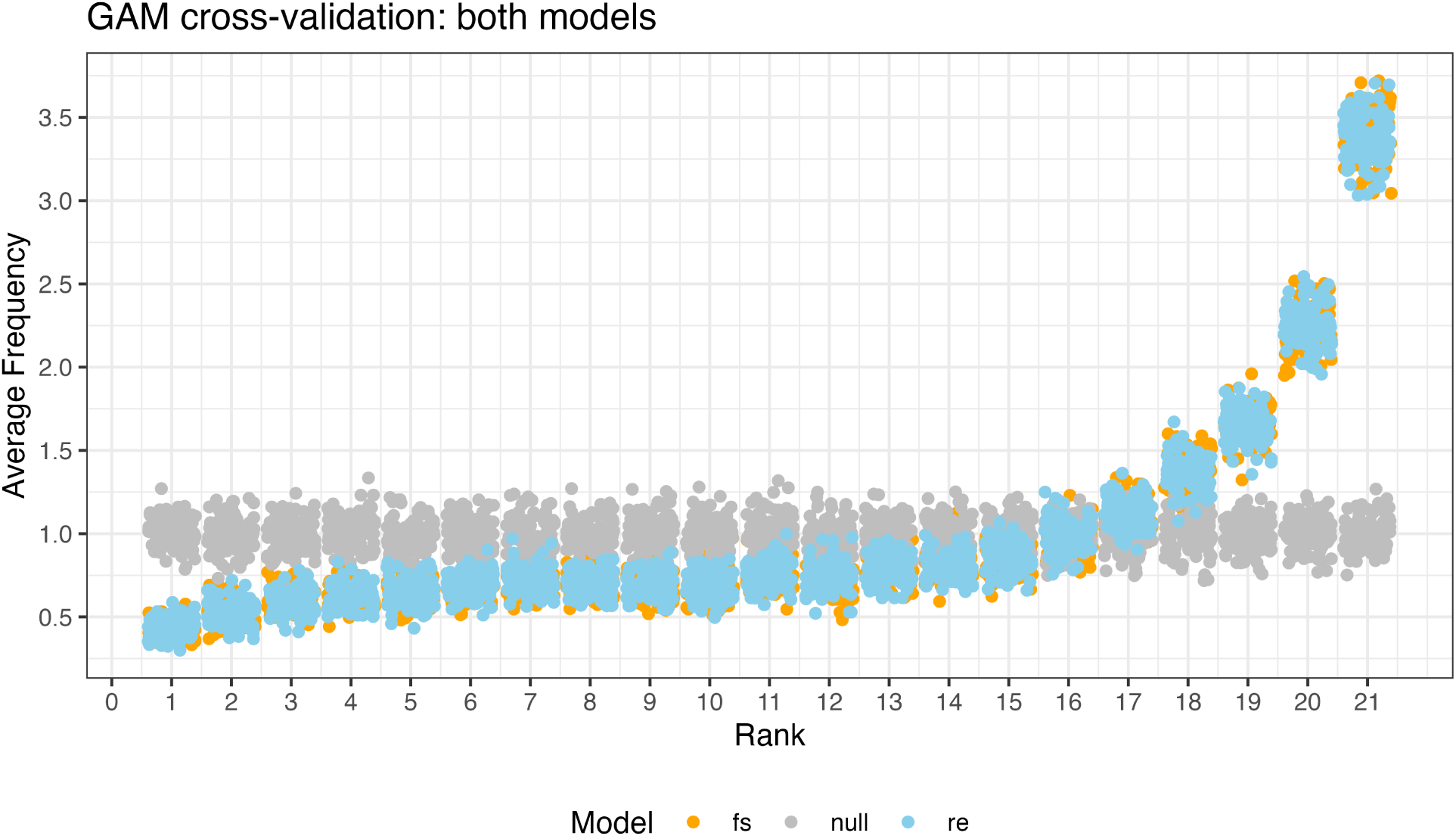
Comparison of cross-validation results for both the final and alternative models. Both models showed very similar performance, suggesting that the model estimates were robust to the nature of the individual-level random effect structure. ‘fs’ is the model with a factor smooth random effect structure, ‘re’ is the random intercept, and ‘null’ is a null model where the rank was selected randomly from the observed and proposed steps.

### **D.** Space use

#### D.1. Minimum convex polygon space use

**Figure S12:**
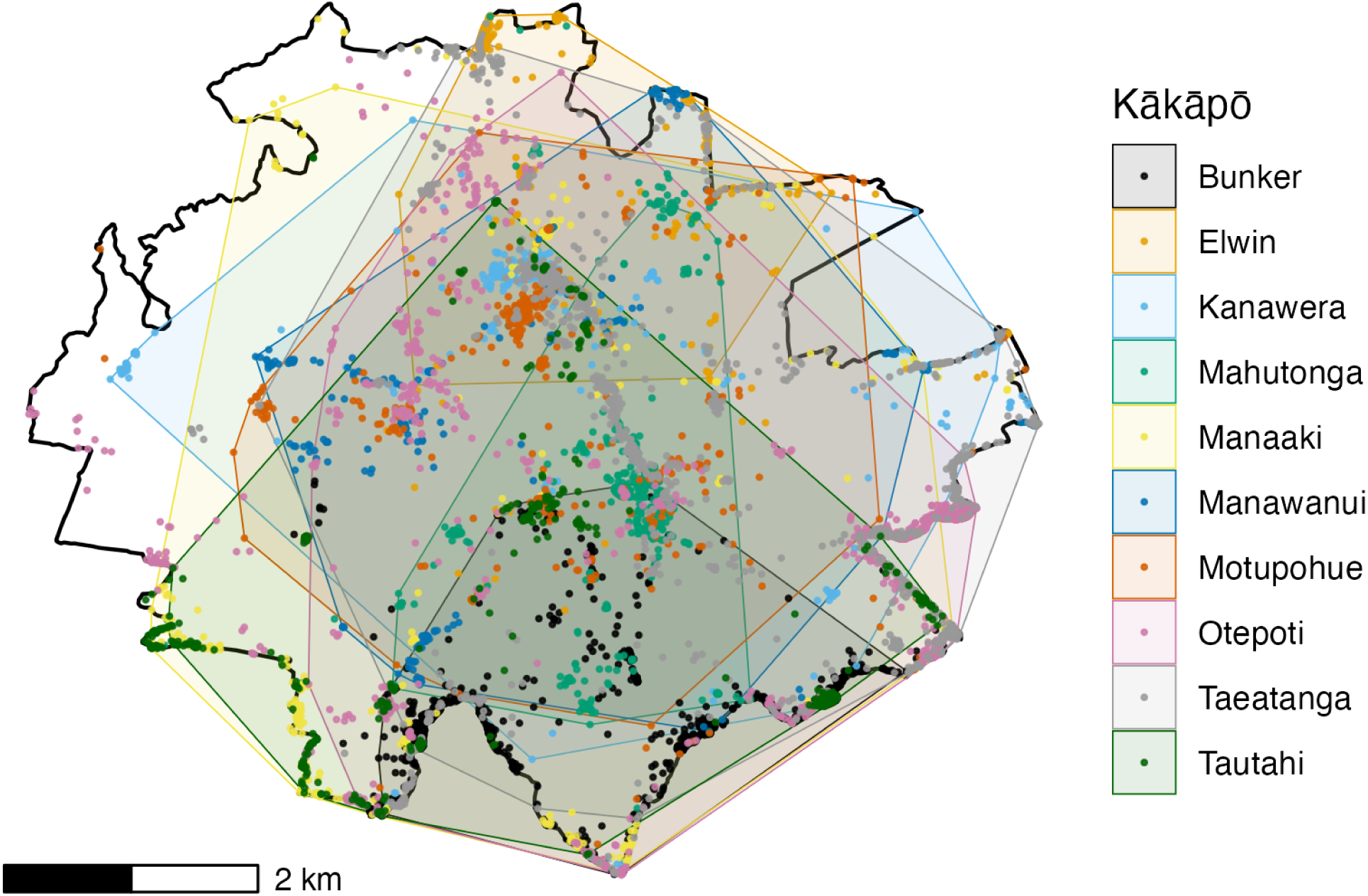
The 95% minimum convex polygons (MCP95) for each kākāpō (shaded areas) derived over all time periods from the locations input to the SSF model (filled circles).

#### D.2. Occurrence Distributions for each kākāpō

The following figures show the occurrence distributions (blue line = 95th percentile, green = 90th percentile, red = 50th percentile) and individual fixes (black circles) for each time step for each kākāpō.

**Figure S13:**
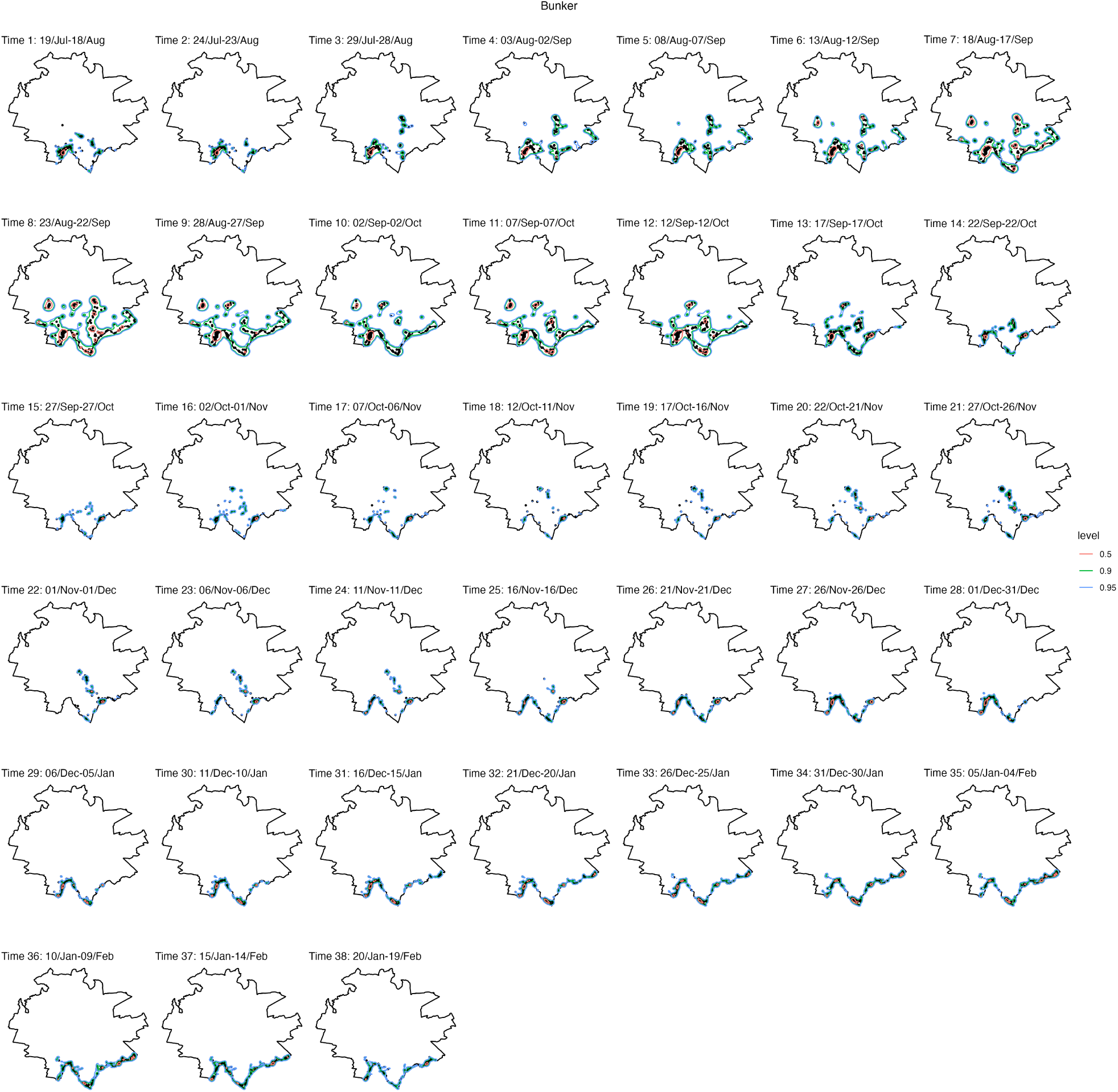
Bunker occurrence distributions and observed locations.

**Figure S14:**
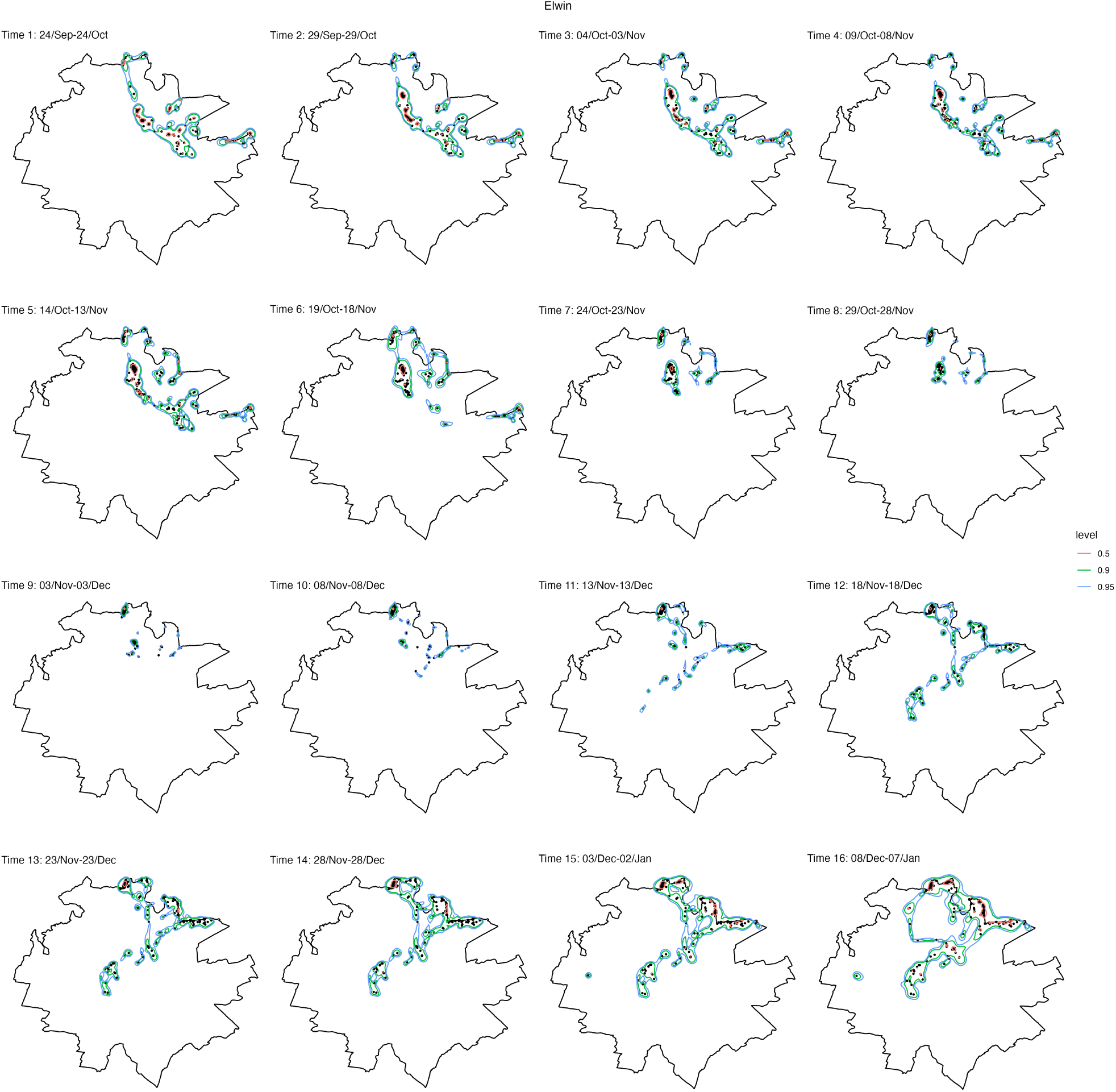
Elwin occurrence distributions and observed locations.

**Figure S15:**
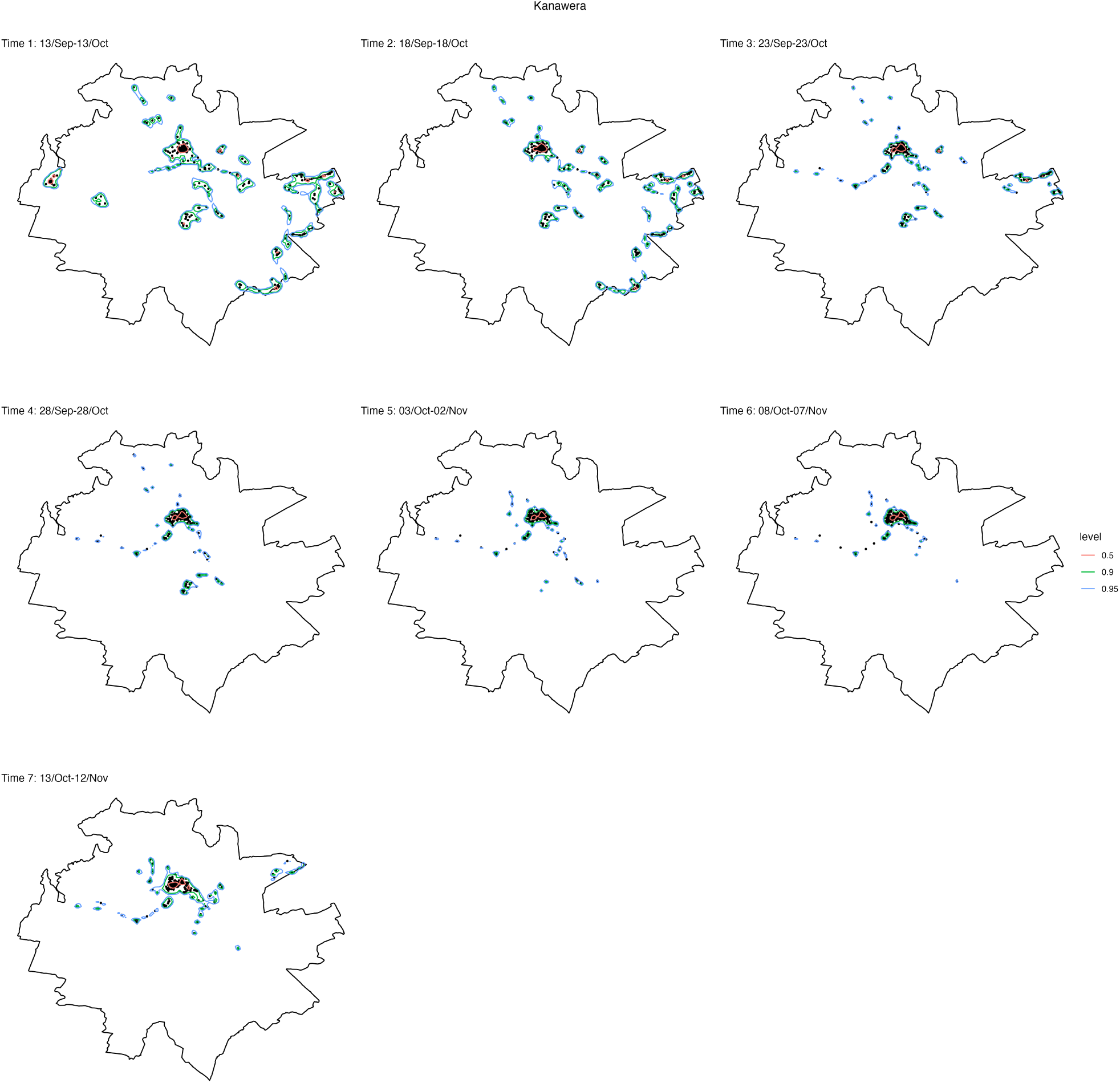
Kanawera occurrence distributions and observed locations.

**Figure S16:**
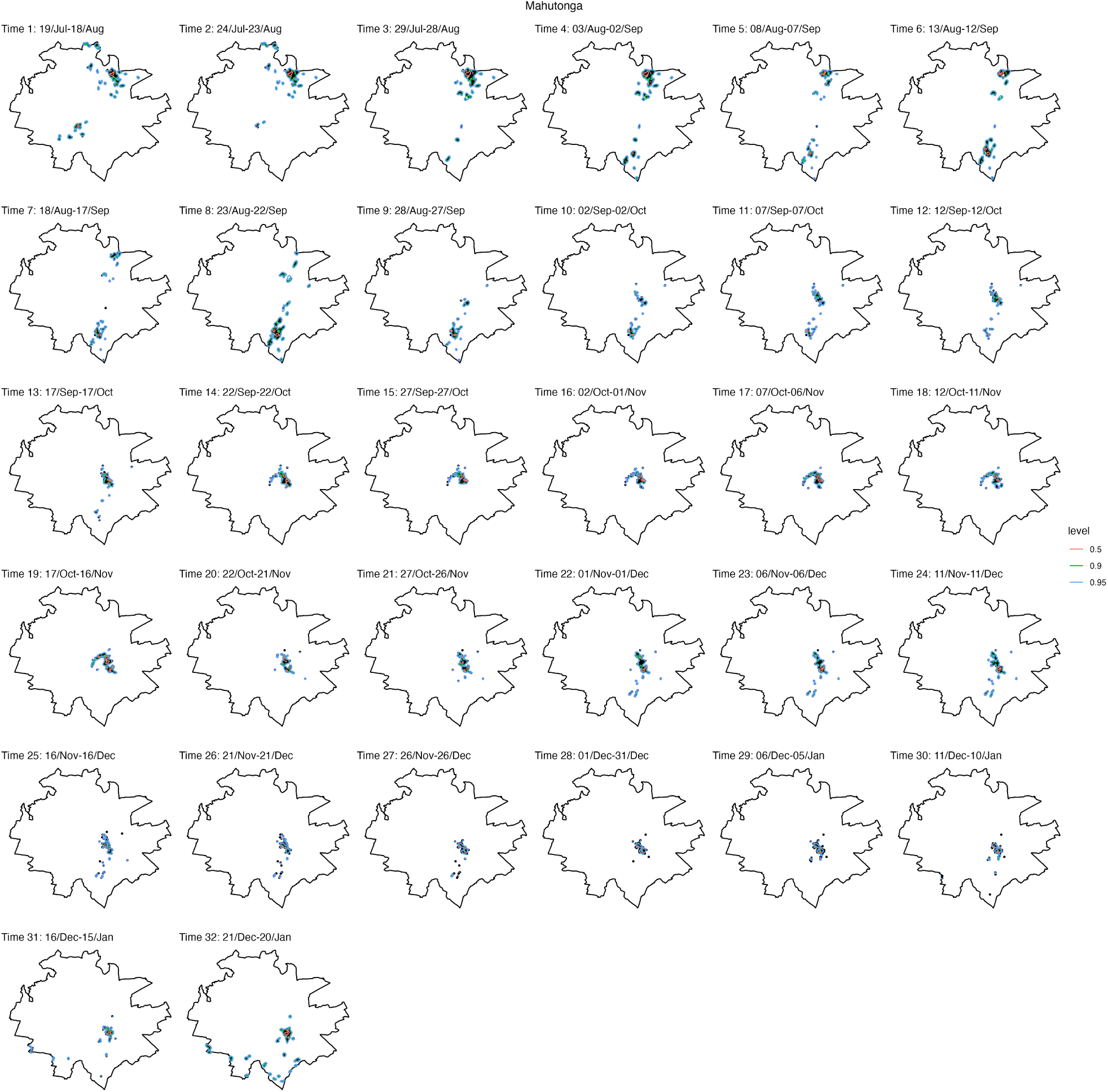
Mahutonga occurrence distributions and observed locations.

**Figure S17:**
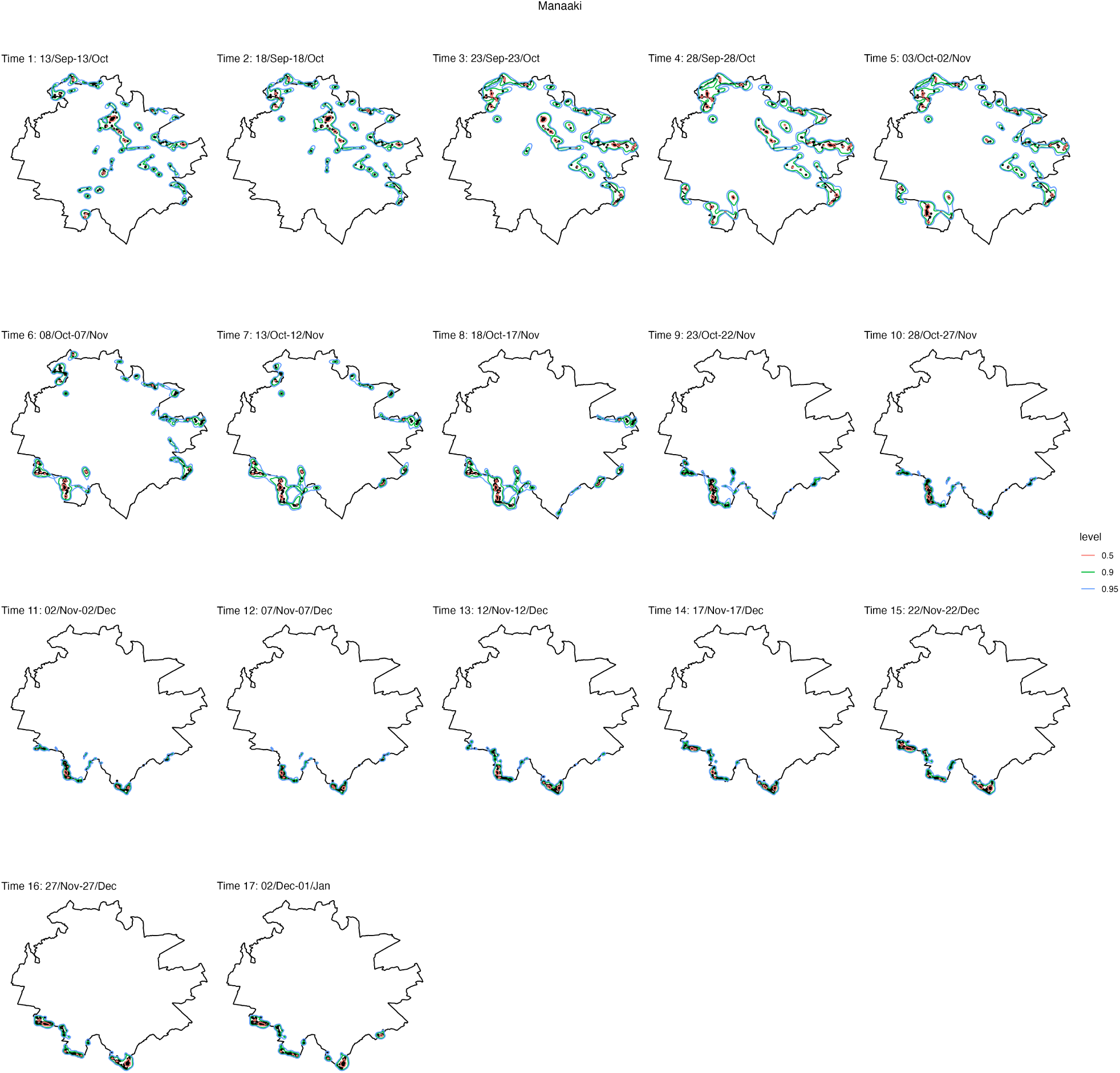
Manaaki occurrence distributions and observed locations.

**Figure S18:**
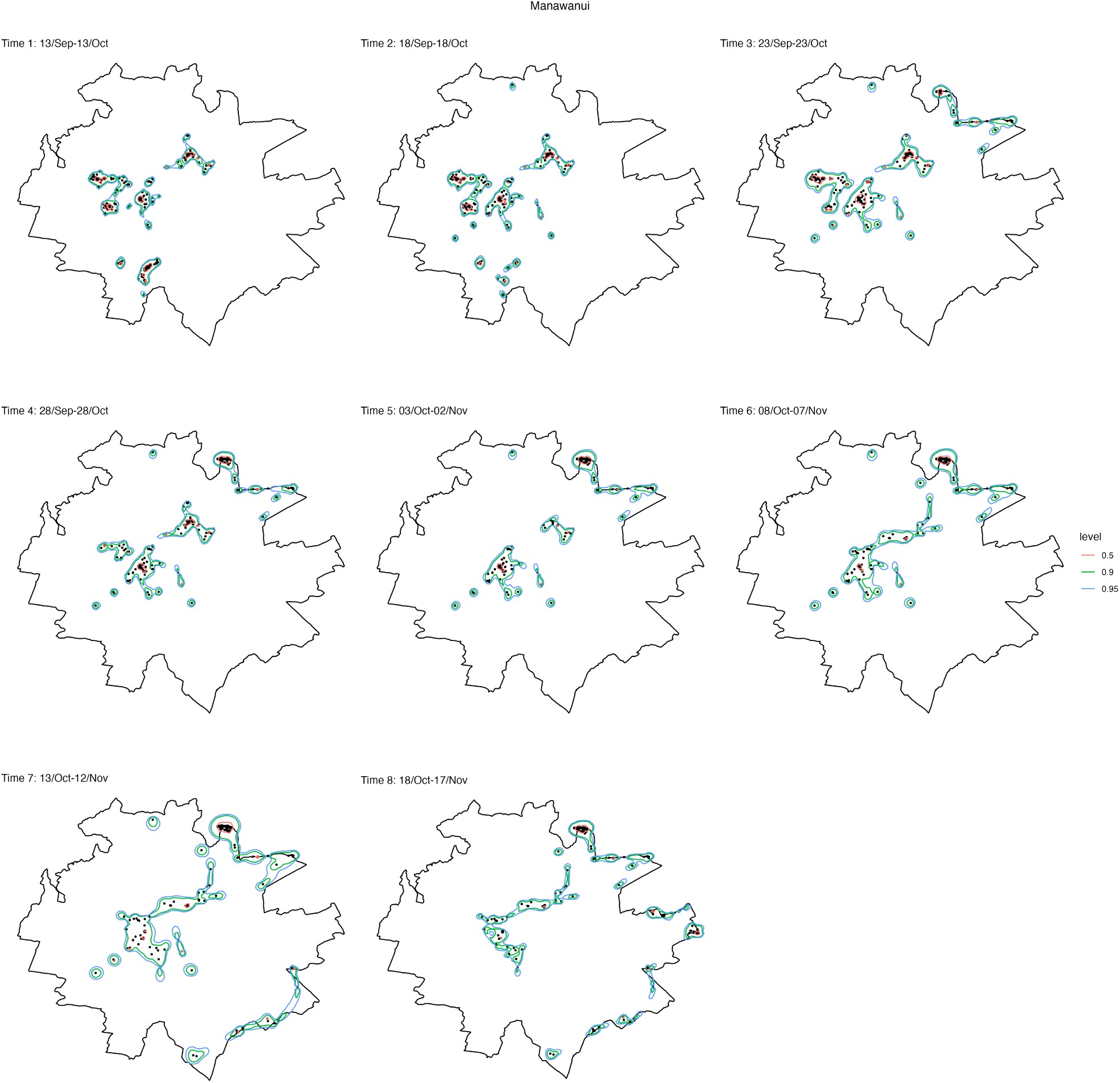
Manawanui occurrence distributions and observed locations.

**Figure S19:**
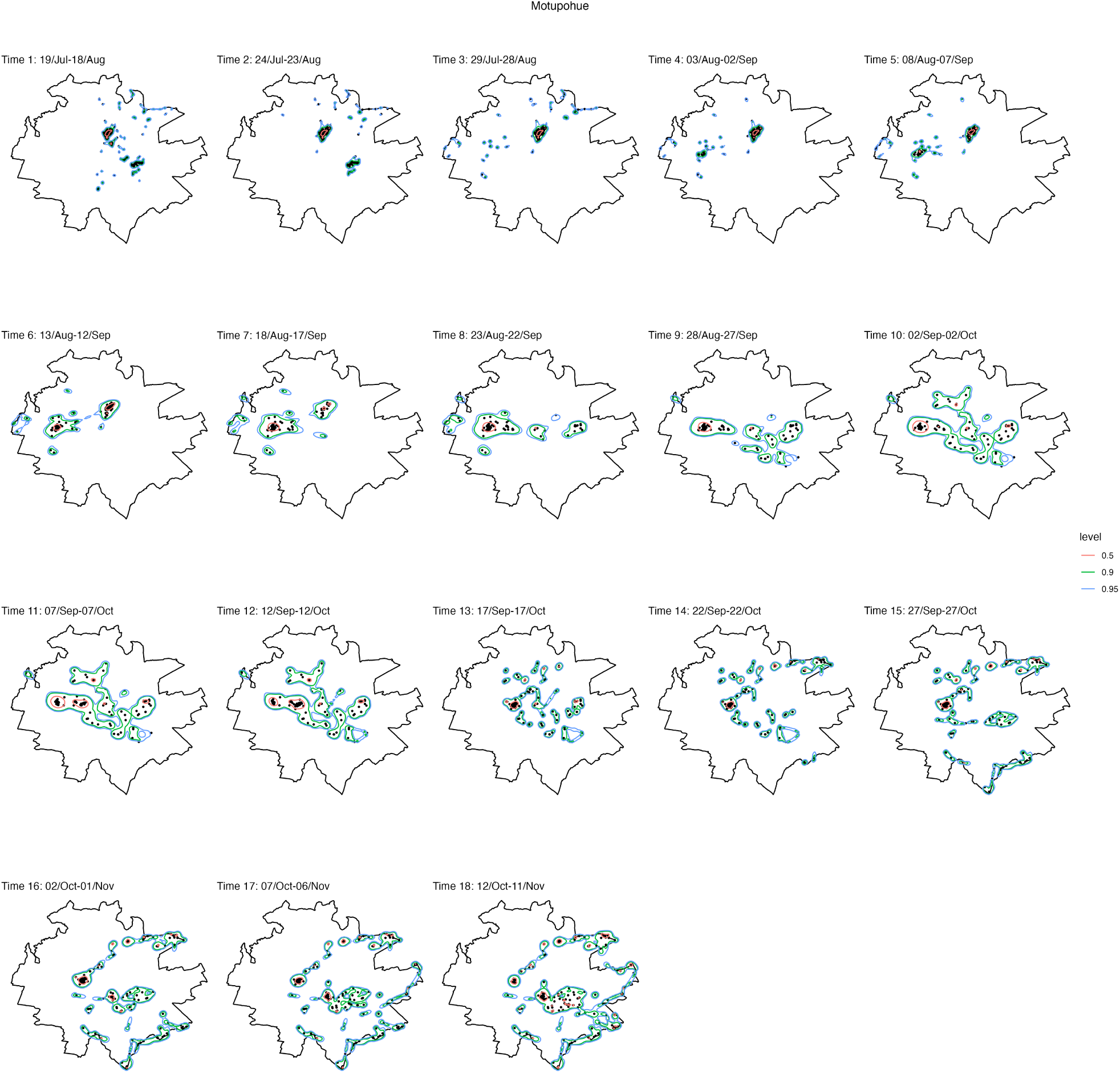
Motupohue occurrence distributions and observed locations.

**Figure S20:**
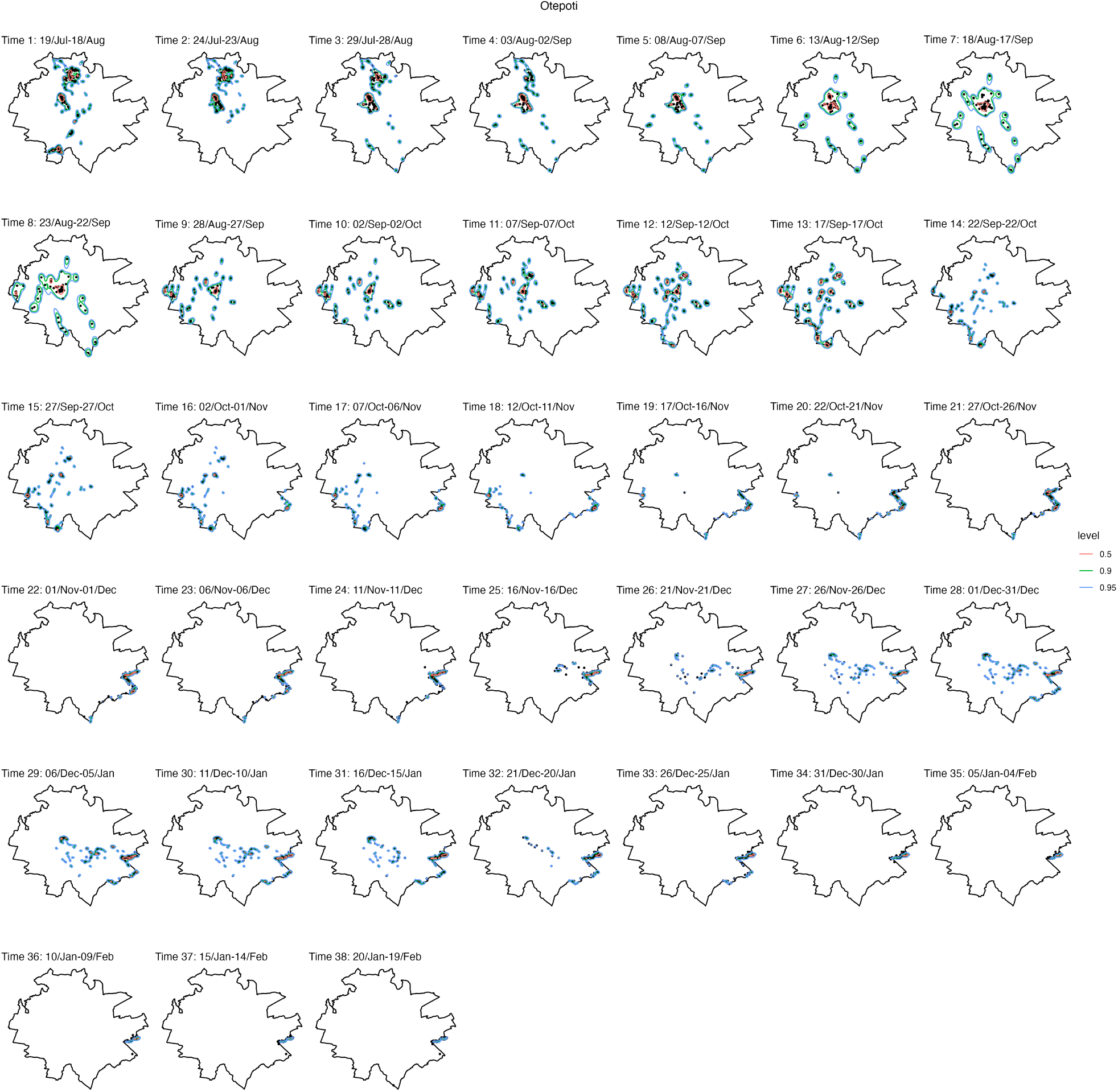
Otepoti occurrence distributions and observed locations.

**Figure S21:**
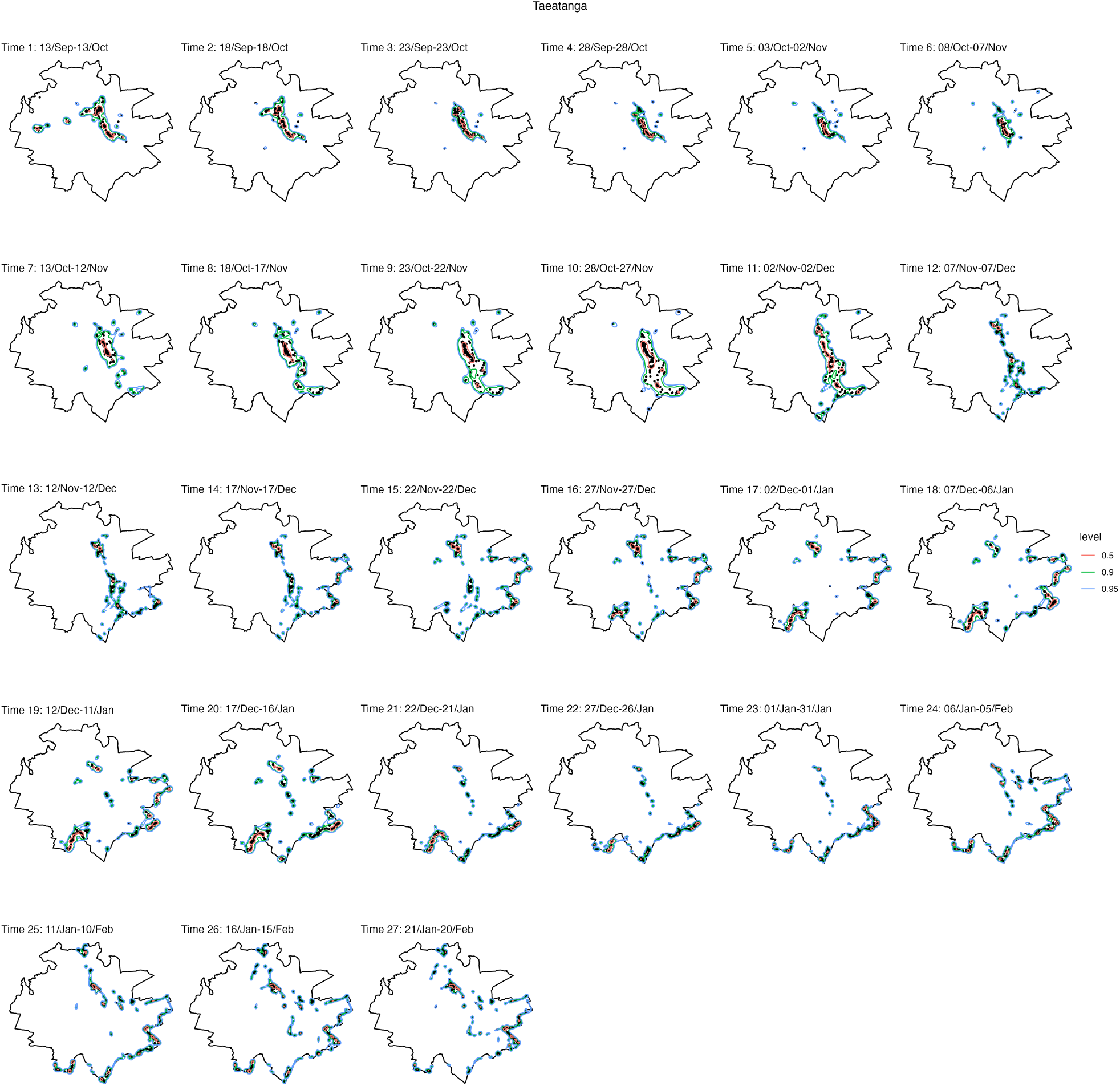
Taeatanga occurrence distributions and observed locations.

**Figure S22:**
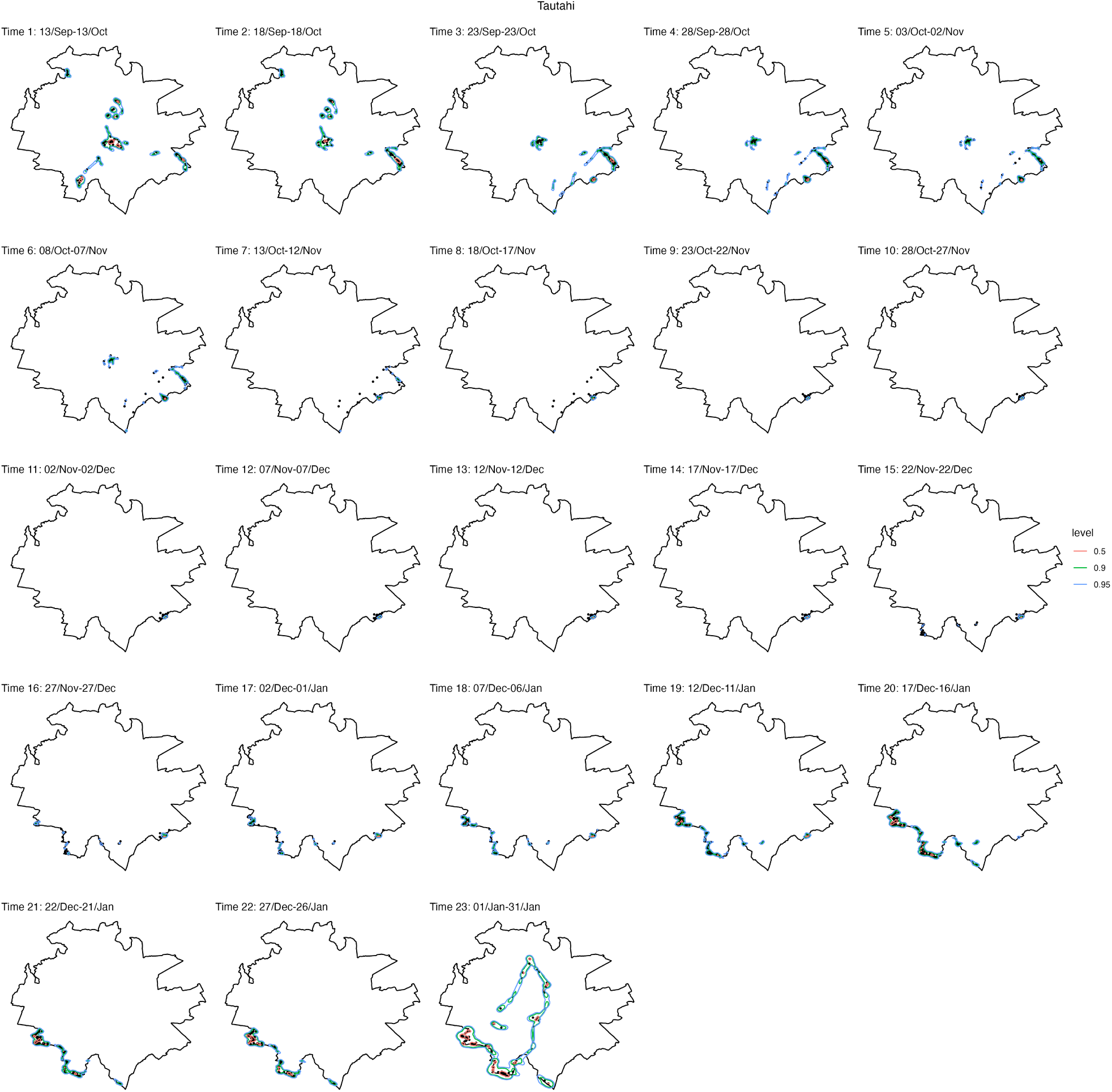
Tautahi occurrence distributions and observed locations.

## Notes

### Competing Interest Statement

The authors have declared no competing interest.

